# Capping protein regulates the balance of assembly among diverse actin networks in *C. elegans* zygotes

**DOI:** 10.64898/2026.03.23.713757

**Authors:** Sarah E. Yde, Cristian Suarez, Shinjini Ray, Ronen Zaidel-Bar, Rachel S. Kadzik, Ed Munro, David R. Kovar

## Abstract

Actin cytoskeleton networks exhibit specialized architectural properties for specific cellular tasks, as determined by the actin-binding proteins (ABPs) associated with each network. Proper allocation of a limiting pool of actin monomers also helps shape the assembly of different F-actin networks. The ABP capping protein (CP) modulates F-actin network architecture through regulation of actin filament length by capping filament barbed ends. Using a combination of *in vitro* biochemistry and quantitative live-cell imaging, we characterize CP as a major regulator of inter-network competition between filopodia and mini-comets, two F-actin networks in the one-cell *C. elegans* embryo (zygote). We establish that this regulation is facilitated in part by competition for binding barbed ends between CP and the F-actin elongator formin CYK-1. Together, these results reveal a role for CP in determining F-actin network architecture and dynamics, regulating the coordination between actin assembly factors to assemble and maintain different dynamic F-actin networks, and allocation of G-actin between competing cortical F-actin networks.

**Summary for table of contents:** Cells assemble diverse actin cytoskeleton networks within a common cytoplasm for essential cellular processes. Yde et al. establish a role for Capping Protein, a regulator of actin filament length, in coordinating the balanced assembly of distinct actin networks in the *C. elegans* zygote.

## Introduction

Cells assemble diverse filamentous (F-) actin networks from a cytoplasmic pool of unassembled globular (G-) actin to facilitate a variety of fundamental cellular processes, including polarization, migration, cytokinesis, and transport of cellular materials (Blanchoin et al. 2014; Pollard and Cooper 2009; Pollard and O’Shaughnessy 2019). The architecture and dynamics of each network are tuned specifically to its cellular roles, and cells must coordinate the assembly of multiple distinct F-actin networks in space and time to serve these different functions. A common pool of cytoplasmic actin and actin binding proteins (ABPs) provides a cell with the versatility to address cellular needs that change over time. Different subsets of ABPs provide F-actin networks with specificity by tuning a network’s architecture and dynamics, and through these, its mechanical properties. Thus, a fundamental challenge is to understand how cells coordinate the self-organization of a shared pool of G-actin monomers and ABPs into multiple distinct F-actin networks with the correct architectures and spatiotemporal dynamics.

F-actin networks are dynamic, in a constant state of assembly and disassembly. Furthermore, self-sorting of ABPs influences the architecture and dynamics of each network (Kadzik et al. 2020), while competition among networks for G-actin limits the size and dynamics of coexisting networks (Suarez et al. 2015; Burke et al. 2014; Billault-Chaumartin and Martin 2019; Suarez and Kovar 2016; Antkowiak et al. 2019). It is therefore critical to elucidate the molecular mechanisms underlying these competitive/homeostatic interactions, and how they affect the characteristics of multiple distinct F-actin networks sharing a common cytoplasm.

One key regulator of F-actin network architecture and actin monomer distribution is capping protein (CP, mammalian CapZ). CP is a heterodimer of two structurally similar subunits (α and β subunits in mammals, CAP-1 and CAP-2 in *C. elegans*) that binds with high affinity (subnanomolar K_D_ in mammals) to the fast-growing barbed ends of actin filaments, preventing addition or loss of actin subunits (Isenberg et al. 1980; Wanger and Wegner 1985; Caldwell et al. 1989; Cooper et al. 1984; Cooper and Pollard 1985). High CP activity near newly nucleated F-actin favors shorter filaments, which are stiffer and better-suited to support protrusive forces that act, for example, at the leading edge of a migrating cell. Lower CP activity allows for longer filaments, which are typically bundled, and better-suited to support long-range transport or the generation of contractile forces by myosin II, as in the contractile ring of a dividing cell (Pollard and Borisy 2003; T. M. Svitkina and Borisy 1999; Mogilner and Oster 1996; Muresan et al. 2022). Thus, by regulating actin filament length, CP tunes the mechanical properties of different F-actin networks (Kasza et al. 2010).

CP also plays distinct roles in controlling the rates of branched F-actin network assembly by the Arp2/3 complex and linear filament assembly by formin or Ena/VASP. CP promotes Arp2/3 complex-mediated branched actin assembly via multiple mechanisms, including barbed end interference and monomer funneling. Barbed end interference is a process whereby CP prevents sequestration of Arp2/3 complex activators (Funk et al. 2021). Monomer funneling is a process whereby CP limits the number of assembly-competent barbed ends, thereby increasing the G-actin pool for productive barbed ends to assemble and promote cell migration (Carlier and Pantaloni 1997; Shekhar and Carlier 2017; Funk et al. 2021; Akin and Mullins 2008). CP also directly competes with the processive F-actin elongators formin and Ena/VASP for barbed ends, thus antagonizing linear F-actin assembly (Shekhar et al. 2015; Bombardier et al. 2015; Harris et al. 2004; Zigmond et al. 2003; Barzik et al. 2005; Bear et al. 2002; Moseley et al. 2004; Winkelman et al. 2014).

Because CP promotes Arp2/3 complex-while antagonizing formin– and EnaVASP-mediated actin assembly, perturbing CP function in cells affects the assembly of branched and linear F-actin networks in different ways. For example, CP perturbation (knockdown, knockout, or mutation) phenotypes include a decreased abundance of branched lamellipodial networks, increased abundance of filopodia, slowing of network assembly dynamics, and hypercontractility phenotypes associated with increased linear actin assembly (Ray et al. 2023; Fan et al. 2011; Jo et al. 2015; Sinnar et al. 2014; Hug et al. 1995; Amatruda et al. 1990; Schafer et al. 1995; Fernández et al. 2011; Hopmann and Miller 2003; Kovar et al. 2005; Terry et al. 2018; Mejillano et al. 2004). While these studies have emphasized roles for CP in the assembly of individual F-actin networks, the distinct effects on different F-actin networks within a cell implies that CP could also regulate the balance of assembly among different F-actin networks that coexist within a common cytoplasm (Billault-Chaumartin and Martin 2019). However, this hypothesis has not been systematically tested in an animal model.

The one-cell *C. elegans* embryo (a.k.a. the zygote) provides a powerful context in which to study the role of CP in coordinating the assembly, lifetime and organization of multiple actin cytoskeleton networks within a common cytoplasm (Fig. 1 A) (Hirani et al. 2019; Chan et al. 2019; Velarde et al. 2007; Shivas and Skop 2012; Skop 2004; Li and Munro 2021; Strome 1986). The zygote is an ideal system for studying the mechanisms of F-actin network self-organization because it is a large cell that assembles multiple dynamic networks throughout its lifetime, is genetically tractable, and is amenable to high spatiotemporal resolution time-lapse fluorescence microscopy, including single molecule quantitative imaging.

**Figure 1.**
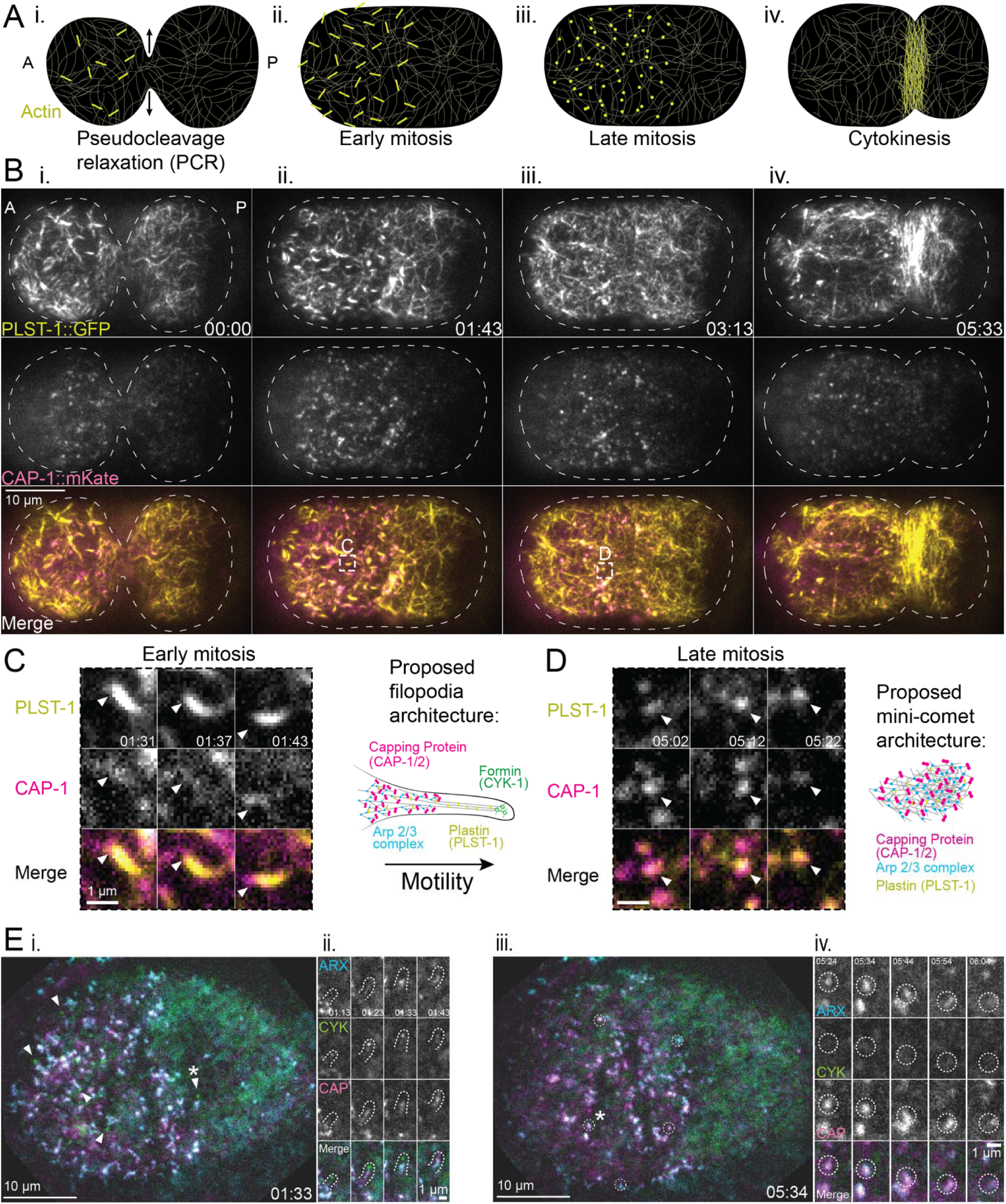
*Ce*CP localizes to the major cortical F-actin networks in the *C. elegans* zygote, including filopodia and mini-comets. (**Ai-iv**) Schematic overview of F-actin networks observed in the *C. elegans* zygote during the first cell cycle. (A, anterior; P, posterior in all panels). A cortical meshwork of formin CYK-1-mediated bundles (thin curved lines) is present at all stages i-iv. Filopodia (thick lines) appear during late interphase (i) and persist through early mitosis (ii). Mini-comets (bright spots) are enriched on the anterior cortex in mid-late mitosis (iii)The contractile ring (dense lines) assembles at the equator during cytokinesis (iv). Vertical arrows in (i) indicate pseudocleavage furrow relaxation which we use as a temporal reference (zero timepoint) in B-E. **(B)** Representative live near-TIRFM images of a zygote expressing endogenously tagged PLST-1::GFP and *Ce*CP CAP-1::mKate at the timepoints described in A. Timestamps indicate time after onset of pseudocleavage relaxation (PCR). **(C)** Magnified view of the indicated filopodium from Bii. Arrowhead indicates local enrichment of *Ce*CP. Right, proposed organization of filopodia with *Ce*CP at the base. Arrow indicates direction of filopodia motility. **(D)** Magnified view of the indicated mini-comet from Biii. Right, proposed organization of mini-comets with *Ce*CP throughout. **(E)** Representative near-TIRFM three-color images of a zygote expressing endogenously tagged formin CYK-1::GFP; Arp2/3 complex subunit ARX-7::HALO, and *Ce*CP CAP-1::mKate, reveals the localization of actin assembly factors and *Ce*CP in filopodia and mini-comets. Timestamps correspond to time post-pseudocleavage relaxation. i. An embryo in early mitosis. Arrowheads indicate representative filopodia. ii. Magnified view of the filopodium indicated by asterisk in i. iii. The same embryo in late mitosis. Dashed circles outline representative mini-comets. iv. Magnified view of the mini-comet indicated by the asterisk in iii.

Throughout the first cell cycle, the zygote assembles multiple F-actin networks at specific times and places to orchestrate polarization and asymmetric cell division (Fig. 1 A) (Hill and Strome 1988). During interphase (lasting ∼10 min), a cortical actomyosin network assembled by the formin CYK-1 and nonmuscle myosin II (NMY-2) generates cortical flows that transport polarity determinants to distinct anterior and posterior cortical domains, divided by a transient pseudocleavage furrow (Fig. 1 A i) (Munro et al. 2004). This network, which we refer to here as the cortical meshwork, also contributes to maintaining normal cortical polarity through mitosis (lasting ∼6 min). Subsequently, during anaphase, formin assembles linear filaments to build and constrict an equatorial contractile ring and complete cytokinesis (Fig. 1 A iv) (Maddox et al. 2005; Swan et al. 1998; Davies et al. 2014).

In addition to these contractile networks, the polarized zygote assembles two additional F-actin structures, which we refer to here as filopodia and mini-comets (Fig. 1 A ii-iii) (Scholze et al. 2018; Hirani et al. 2019; Shivas and Skop 2012). Filopodia are membrane-bound actin filament bundles with formin at their tips (Hirani et al. 2019), which are enriched on the anterior cortex from late interphase into early mitosis (Fig. 1 A ii, Fig. S1 A, yellow). Although their function in the zygote is unclear, similar structures extend between adjacent blastomeres during later embryonic stages and may therefore play a role in intercellular contact formation and/or signaling, as with canonical filopodia found in numerous cell types (Mattila and Lappalainen 2008). Mini-comets are endocytic structures assembled by the Arp2/3 complex that are enriched in the anterior cortex during mid-late mitosis and are thought to play a role in maintaining correctly-sized polarity domains (Fig. 1 A iii, Fig. S1 A, magenta) (Shivas and Skop 2012; Nakayama et al. 2009). Filopodia and mini-comets overlap with the formin-mediated cortical F-actin meshwork, and they harness different modes of actin assembly, providing a unique opportunity to explore the mechanisms that coordinate the assembly of multiple F-actin networks within a common cytoplasm, and specifically to evaluate the contributions of CP to this complex coordination.

Here, using a combination of *in vitro* biochemistry and quantitative time-lapse microscopy of both control and CP-depleted zygotes, we found that the assembly dynamics of filopodia and mini-comets are in balance with one another, that CP activity is an important regulator of this balance, and that competition between CP and formin is a major mechanism of this regulation. Thus, maintaining a proper balance between CP and formin activity has significant roles in regulating the allocation of actin to different F-actin networks. This study moves us closer to a molecular understanding of how ABPs can have diverse effects on different F-actin networks and thereby contribute to self-organization of distinct actin cytoskeleton networks.

## Results

### Capping protein localizes to multiple cortical F-actin networks in the *C. elegans* zygote

*C. elegans* Capping Protein (*Ce*CP) CAP-1/CAP-2 was shown to be important for *C. elegans* gonad development and spermatheca contractility (Ray et al. 2023; Saini et al. 2025), but its roles in controlling the assembly of specific F-actin networks was not determined at the levels of single cells, F-actin networks, or single molecules. To understand how CP contributes to actin cytoskeletal architecture and dynamics in the zygote, we examined the localization of *Ce*CP to distinct F-actin networks throughout the first mitotic cell cycle in *C. elegans* zygotes. We used near-TIRF microscopy (near-TIRFM) to image zygotes co-expressing endogenously tagged *Ce*CP (CAP-1::mKate (Ray et al. 2023)), and endogenously-tagged plastin as a general marker for F-actin networks (PLST-1::GFP (Ding et al. 2017)) (Fig. 1 B, Video 1).

We found that CAP-1::mKate localizes to all four cortical F-actin networks marked by PLST-1::GFP: the cortical meshwork, filopodia, mini-comets, and the contractile ring (Fig. 1 B, Video 1). Small puncta of CAP-1::mKate are distributed throughout the cortical meshwork at all stages and are slightly enriched ∼1.3-fold in the contractile ring during cytokinesis (Fig. 1 B iv and Fig. S1 B, Video 1). During early and late mitosis, dynamic accumulations of CAP-1::mKate are concentrated in filopodia and mini-comets (Fig. 1 B ii-iii). Consistent with previous results (Hirani et al. 2019), filopodia are enriched on the anterior cortex during late interphase/early mitosis (Fig. S1 A, pseudocleavage relaxation) and mini-comets are enriched in late mitosis (Fig. S1 A, 100 s before contractile ring assembly). In early mitosis (Fig. 1 B ii), filopodia appear as elongated structures that undergo persistent directional motion within the cortical imaging plane (Fig. 1 C, Video 2, upper). CAP-1::mKate localizes to the base of filopodia but is mostly excluded from the shafts and tips (Fig. 1 C, Video 2, upper). Conversely, in late mitosis (Fig. 1 B iii), mini-comets appear as smaller, more punctate structures that undergo local movements without directional persistence (Fig. 1 D, Video 2, lower). Unlike in filopodia, CAP-1::mKate localizes throughout the entire mini-comet (Fig. 1 D, Video 2, lower).

To more fully characterize *Ce*CP localization, we imaged zygotes co-expressing endogenously-tagged CAP-1::mKate with endogenously tagged versions of the two major F-actin assembly factors in the *C. elegans* zygote: formin CYK-1 (CYK1::GFP) and the Arp2/3 complex (tagged with Arp2/3 complex component ARX-7::HALO) (Fig. 1 E, Video 3). Simultaneous imaging of *Ce*CP, CYK-1 and Arp2/3 complex distinguishes filopodia from mini-comets, as CYK-1 puncta mark the tips of filopodia (Fig. 1 E i-ii), whereas mini-comets do not contain CYK-1 (Fig. 1 E iii-iv). In filopodia, dynamic clouds of ARX-7::HALO and CAP-1::mKate follow closely behind, but do not colocalize with individual CYK-1::GFP puncta, indicating that *Ce*CP and Arp2/3 complex dynamically colocalize at the bases of filopodia, but are excluded from the formin-mediated tips (Fig. 1 E ii, Video 4, upper). In mini-comets, CAP-1::mKate colocalizes with ARX-7::HALO throughout the entire mini-comet (Fig. 1 E iii-iv, Video 4, lower). In summary, *Ce*CP localizes to all the major cortical F-actin networks in the *C. elegans* zygote, but with specific localization patterns, indicating the possibility of unique roles for *Ce*CP in organizing and regulating these architecturally diverse F-actin networks.

### The residence time of capping protein on barbed ends differs *in vivo* and *in vitro*

The biochemical properties of capping proteins from a diverse range of species have been well-characterized *in vitro*, revealing that they have a wide range of affinities for F-actin barbed ends, ranging from ∼0.1-100 nM (Kuhn and Pollard 2007; Wear et al. 2003; Schafer et al. 1996; Wirshing et al. 2023; Kim et al. 2012). To determine where *C. elegans* capping protein (*Ce*CP) falls on the spectrum of weak to strong cappers, we expressed and purified recombinant *Ce*CP (CAP-1/CAP-2) from *E. coli*. We directly compared *Ce*CP to the well-characterized mouse CP (*Mm*CP) (Kim et al. 2012) in bulk and single-filament *in vitro* assays (Fig. 2).

**Figure 2.**
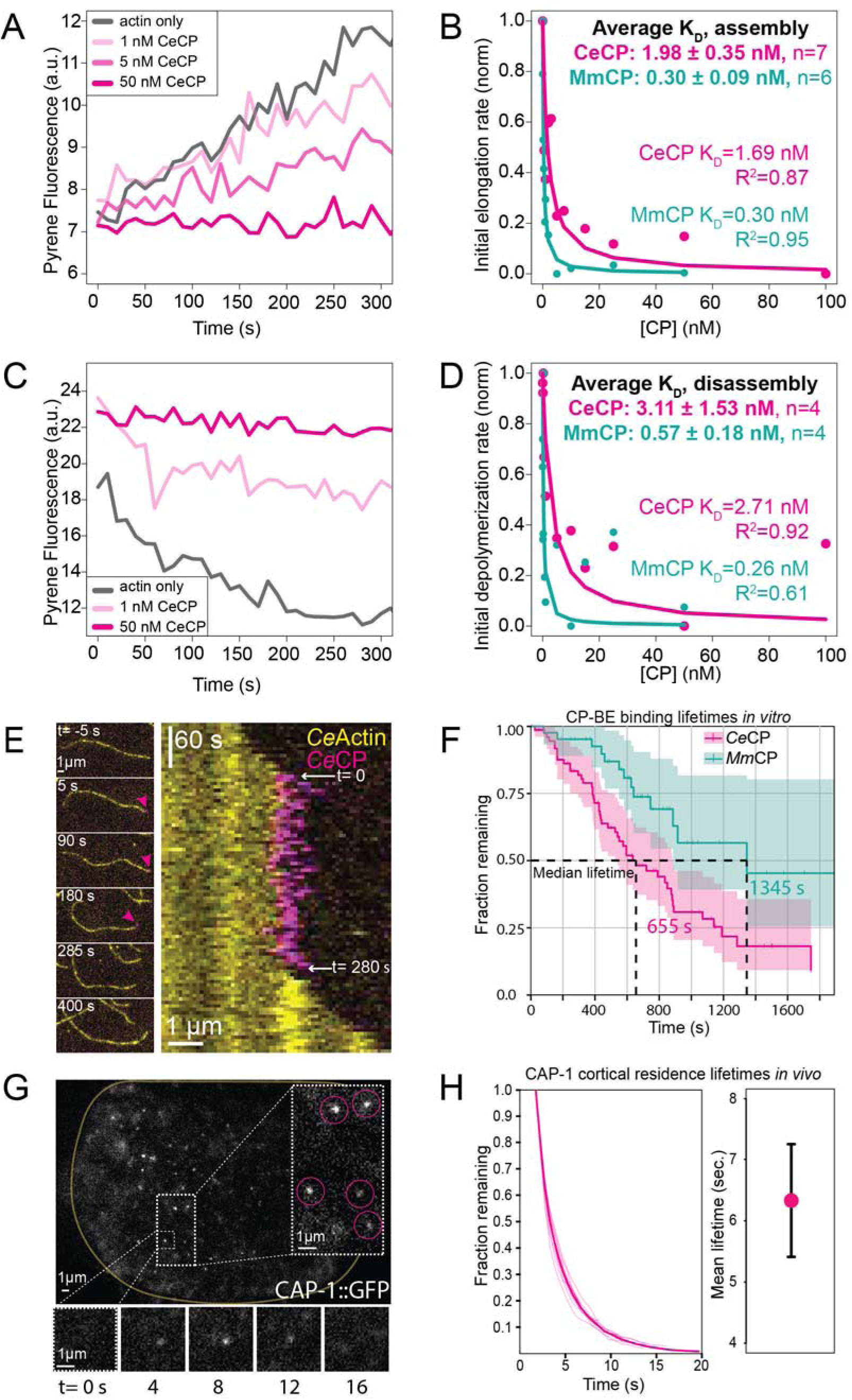
Comparing *Ce*CP binding dynamics *in vitro* and *in vivo* suggests that *Ce*CP stays bound to a barbed end for the entire lifetime of an actin filament. (**A-B**) *In vitro* seeded elongation assay, initiated by addition of 0.5 µM Mg-ATP actin monomers (10% pyrene-labeled) to 0.5 µM pre-assembled unlabeled F-actin seeds. **(A)** Representative time course for actin alone (grey) and for the indicated concentrations of CeCP. **(B)** Dependence of the seeded actin assembly rate (linear slopes from 50-500s) on the concentration of *Ce*CP (magenta) or mouse *Mm*CP (green). Dissociation constants (K_D_) determined via nonlinear regression analysis displayed as: light font (shown replicate), and bold font (average of all replicates). (C-D) *In vitro* actin filament disassembly assay. **(C)** Representative time course of the barbed end depolymerization of 5 µM preassembled actin filaments (50% pyrene-labeled) after dilution to 0.1 µM in the presence of the indicated concentrations of *Ce*CP. **(D)** Dependence of disassembly rate (linear slopes from 50-300s) on the concentration of *Ce*CP (magenta) or *Mm*CP (green). Dissociation constants (K_D_) determined via nonlinear regression analysis are displayed as: light font (shown replicate), and bold font (average of all replicates). **(E-F)** Two-color TIRFM visualization of the assembly of 1.5 µM Mg-ATP *C. elegans* actin (10% alexa488-labeled) with 647-labeled CP. **(E)** (left) Representative time-lapse images of a single *Ce*CP-647 molecule (magenta arrowhead) at the barbed end of an actin filament. (Right) Kymograph of the filament before, during, and after the *Ce*CP binding event. **(F)** Kaplan-Meyer analysis of barbed end binding event lifetimes for *Ce*CP (magenta) and *Mm*CP (green). Dashed vertical lines indicate median survival times, p=0.0024. Shaded areas represent 95% confidence intervals. **(G-H)** Single molecule analysis of CeCP dissociation kinetics *in vivo*. **(G)** Near-TIRFM visualization of single *Ce*CP particles at the cortex of a *C. elegans* zygote (yellow outline) overexpressing CAP-1::GFP (*Ce*CP) at low levels. Dashed box at right shows magnified view of single *Ce*CP particles (magenta circles) detected by our analysis. (Lower boxes show magnified views of a single tracked CAP-1::GFP particle over time. **(H)** (Left) Decay curves for single CAP-1 particles at the cortex of *C. elegans* zygotes show the fraction of single molecules (fraction remaining) with lifetimes > time t as a function of t. Faint lines show data from individual embryos; opaque line shows data pooled over N = 9 embryos. (Right) Mean residence times estimated from exponential fits to the pooled decay curve, with correction for photobleaching. Error bars indicate 95% confidence intervals obtained by bootstrap sampling (see Methods for details).

To determine the affinity of *Ce*CP for F-actin barbed ends, we used two distinct assays to measure the dependence of bulk actin assembly and disassembly on *Ce*CP concentration. First, we used a seeded pyrene actin assembly assay to measure the effect of varying *Ce*CP concentration on the addition of pyrene-labeled actin monomers (10%) onto preexisting barbed ends of unlabeled F-actin seeds over time (Fig. 2 A-B). Increasing concentrations of *Ce*CP decrease the seeded assembly of actin filaments (Fig. 2 A-B), indicating that the *Ce*CP caps the barbed ends of actin filaments and prevents their elongation. Second, we performed pyrene actin disassembly assays to quantify the effect of varying *Ce*CP concentrations on the barbed end depolymerization of pre-assembled actin filaments (50% pyrene-labeled) upon their dilution to the barbed end critical concentration of 0.1 µM (Fig. 2 C-D). Increasing amounts of *Ce*CP decrease barbed end disassembly rates of F-actin (Fig. 2 C-D), indicating that *Ce*CP caps barbed ends, preventing both polymerization and depolymerization of F-actin. Nonlinear least squares fits to the dependence of the seeded assembly and disassembly rates on the concentration of CP estimates that the dissociation constant (K_D_) of purified *Ce*CP on F-actin barbed ends is ∼2 nM (Fig. 2 B and D, pink). By comparison, *Mm*CP has a 10-fold higher affinity for barbed ends, with a K_D_ of ∼0.3 nM (Fig. 2 B and D, green), consistent with previous reports (Kuhn and Pollard 2007; Wear et al. 2003).

To directly observe the kinetics of individual CPs associating with F-actin barbed ends *in vitro*, we labeled purified *Ce*CP and *C. elegans* actin and used TIRFM to visualize spontaneous actin assembly in the presence of labeled *Ce*CP. Single molecules of labeled SNAP-647 *Ce*CP bind to growing barbed ends of assembling *C. elegans* actin (Fig. 2 E, Video 5), halting barbed end assembly, until *Ce*CP dissociates from the barbed end and filament elongation resumes (Fig. 2 E, kymograph, Video 5). Thus, like other CPs, *Ce*CP directly and reversibly associates with the F-actin barbed end to inhibit barbed end assembly. To quantify the kinetics of *Ce*CP-barbed end binding events, we measured the lifetimes of these events and plotted them on a Kaplan-Meier survival curve (Fig. 2 F). The median binding lifetime for *Ce*CP *in vitro* is 655 seconds, compared to a lifetime of 1,345 seconds for *Mm*CP (Fig. 2 F). Thus, *Ce*CP has a weaker affinity for barbed ends and a faster off-rate (though still on the order of minutes) than *Mm*CP.

*In vitro* experiments reveal *Ce*CP’s association with F-actin barbed ends in the absence of regulatory proteins. However, cells contain other factors that could regulate the ability of CP to bind barbed ends (Edwards et al. 2014). Therefore, we used *in vivo* single molecule imaging and particle tracking analysis (Robin et al. 2014) to quantify the kinetics of *Ce*CP’s association with cortical F-actin in *C. elegans* zygotes transgenically expressing CAP-1::GFP. We used RNA interference (RNAi) against GFP to specifically deplete CAP-1::GFP to levels that allow detection of single molecules at the zygote cortex using near-TIRFM (Fig. 2 G). We collected single molecule data at 0.5 second intervals and performed particle tracking as previously described (Video 6) (Robin et al. 2014; Li and Munro 2021; Lang et al. 2025). As in our previous work, trajectories with shorter residence times (<= 2 seconds) had much higher mobilities, likely reflecting the transient appearance of cytoplasmic CAP-1::GFP molecules in the cortical imaging plane, while longer trajectories had much lower subdiffusive mobilities consistent with cortical F-actin binding (Fig. S2). Therefore, we focused our analysis on molecules with tracked lifetimes >2 s. We constructed release curves that plot the fraction of CAP-1::GFP molecules with tracked lifetimes >t as a function of t (Fig. 2 H) (Robin et al. 2014). These curves were well-fit by a model for exponential decay (Fig. S2), yielding an apparent dissociation rate which is the sum of the CeCP dissociation rate k_dis_ and a photobleaching rate k_pb_. Comparing fits for data obtained with identical unit exposures, but different exposure duty ratios, allowed us to extract simultaneous estimates of the photobleaching and dissociation rates (see Materials and Methods for details). After correcting for photobleaching, we estimate that the mean cortical residence time of CeCP is 6.33+/-0.92 s (Fig. 2 H). Significantly, the mean cortical residence time of CeCP is two orders of magnitude shorter than the average barbed end residence time we measured for CeCP *in vitro* (Fig. 2 F), but it is similar to the lifetime of cortical F-actin measured in *C. elegans* zygotes using similar single molecule approaches (Robin et al. 2014). This suggests that the rate-limiting step for dissociation of CP from the zygote cortex is actin filament disassembly. In particular, the comparison of bound lifetimes measured *in vitro* and *in vivo* suggest that once bound, CP stays bound to cortical actin filaments for their entire lifetimes in the zygote.

### Capping protein regulates the balance of filopodium and mini-comet assembly

*Ce*CP localizes to filopodia, which are assembled by formin CYK-1 and the Arp2/3 complex, as well as to mini-comets, which are assembled by the Arp2/3 complex (Fig. 1 B-E). *Ce*CP colocalizes with the Arp2/3 complex in both structures, but not with formin. Both filopodia and mini-comets are much longer-lived than the lifetimes of polymerized actin and the durations of *Ce*CP-barbed end binding events *in vivo* (Fig. 2 H, Fig. 5 F), suggesting that these structures are maintained despite continuous turnover of F-actin and *Ce*CP. We therefore suspected that *Ce*CP plays key roles in controlling the relative numbers of, or assembly dynamics within, filopodia and mini-comets.

To elucidate roles of *Ce*CP in the differential regulation of these two networks, we used RNAi to deplete the maternal supply of CAP-1 (*cap-1(RNAi)*) in zygotes expressing the general F-actin marker LifeAct::mCherry, and used near-TIRFM to observe the effects on numbers and dynamics of filopodia and mini-comets (Fig. 3 A, Video 7). We focused our analysis during mitosis, when filopodia (early mitosis) or mini-comets (late mitosis) are most abundant. As with PLST-1::GFP (Fig. 1), in zygotes expressing LifeAct::mCherry, filopodia appear as elongated thick bundles of F-actin with characteristic directional motion (Fig. 3 A, outlined in yellow in insets; Video 8, left), while mini-comets appear as small puncta that move with lower directional persistence (Fig. 3 A, outlined in magenta in the insets, Video 8, right).

**Figure 3.**
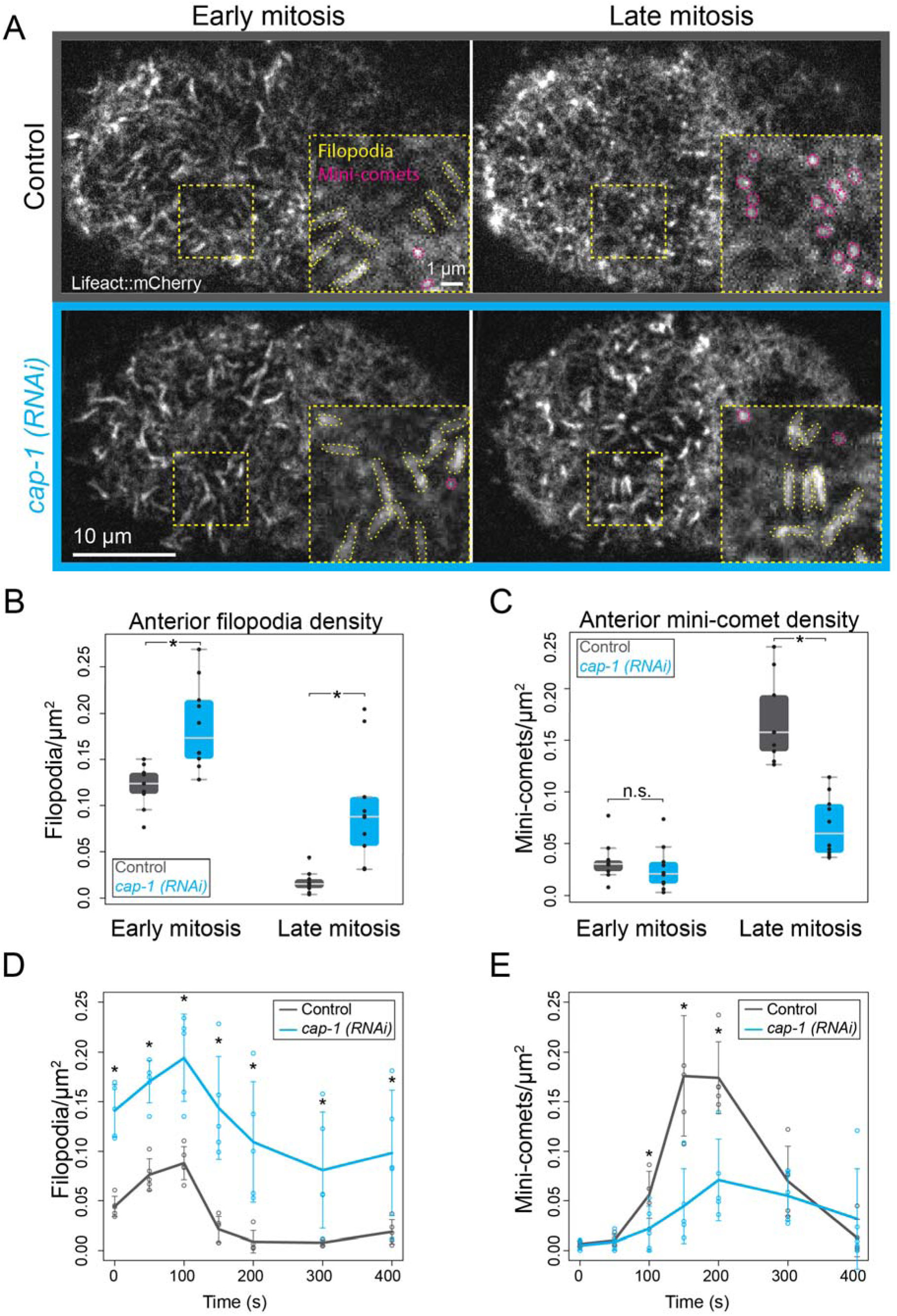
Depletion of *Ce*CP in the zygote increases filopodium abundance and decreases mini-comet abundance. (**A**) Representative images of control (top) and *cap-1*(RNAi) (bottom) embryos during early (left) and late (right) mitosis expressing Lifeact::mCherry and visualized by near-TIRFM. Insets (yellow boxes) of the indicated regions show magnified views of filopodia (dashed yellow lines) and mini-comets (dashed magenta circles). **(B)** Measurements of filopodium density on the anterior cortex during early and late mitosis in control (gray, N=9 embryos) and *cap-1(RNAi)* (blue, N=10 embryos). Black dots represent single embryos. *(early mitosis p=0.0016, late mitosis p=0.0011, unpaired t-tests) **(C)** Mini-comet density on the anterior cortex during early and late mitosis in control (gray, N=9 embryos) and *cap-1 (RNAi)* embryos (blue, N=10). Black dots represent single embryos. * (early mitosis p=0.4146, late mitosis p<0.0001, unpaired t-test). **(D-E)** Filopodium (D) and mini-comet **(E)** density over time in the anterior of control (gray) and *cap-1 (RNAi)* embryos (blue), where time 0 is pseudocleavage relaxation. N=5 embryos per condition. *p<0.05, unpaired t-test.

*Ce*CP depletion appears to induce more filopodia and fewer mini-comets during mitosis compared to controls (Fig. 3 A). To assess this possibility, we measured the density (number of structures per unit area) of filopodia and mini-comets on the anterior cortex at two timepoints during early and late mitosis, 80 seconds and 210 seconds after pseudocleavage furrow relaxation (PCR), respectively (Fig. 3 B-C). Compared to controls, *cap-1(RNAi)* increases the density of filopodia by ∼1.6-fold during early mitosis, and by ∼5.6-fold during late mitosis when control cells have very few filopodia (Fig. 3 B). We confirmed that, like wild-type filopodia, these ectopic filopodia are membrane-bound structures with formin (CYK-1) at the tips and Arp2/3 complex (ARX-7) at the base (Fig. S3, Video 9). Similar to controls, the density of mini-comets remains low during early mitosis in *cap-1(RNAi)* embryos (Fig. 3 C). Conversely, during late mitosis, *cap-1(RNAi)*-treated cells have 2.5-fold lower mini-comet density (Fig. 3 C). Thus, decreasing the amount of *Ce*CP in the zygote increases the density of filopodia, and decreases the density of mini-comets.

To better understand how changes in filopodia and mini-comets are coordinated during mitosis in control and *cap-1(RNAi)* zygotes, we measured filopodia and mini-comet densities at additional timepoints, up to 400 seconds after PCR (Fig. 3 D-E). In control embryos, filopodium density peaks at around 100 seconds after PCR, then decreases throughout late mitosis. In contrast, mini-comet density begins to rise at 100 seconds after PCR, peaks around 200 seconds after PCR, and decreases before cytokinesis. Upon *Ce*CP depletion, the shapes of the curves are similar, but their amplitudes change significantly. In *cap-1(RNAi)* zygotes, filopodium density is significantly increased (∼2-12-fold) at every timepoint (Fig. 3 D), while mini-comet density is significantly reduced (∼2-to 4-fold) throughout its peak assembly timeframe (Fig. 3 E).

Because F-actin network assembly is coupled to the cell cycle, one possible mechanism for increased filopodia at later timepoints could be that the cell cycle progression is perturbed in *cap-1(RNAi)* embryos. To assess this possibility, we compared the timing of cell cycle progression in control and *cap-1(RNAi)* embryos and found that *cap-1(RNAi)* does not have significant effects on cell cycle timing (Fig. S4). Consistent with this observation, the timing with which filopodia and mini-comet assembly rises and falls is preserved in *cap-1(RNAi),* indicating that temporal control over assembly of these networks is still coupled to the cell cycle (Fig. 3 D-E).

Together, these results suggest two major conclusions. First, filopodium and mini-comet assembly are in dynamic balance throughout mitosis, such that increased assembly of one network leads to decreased assembly of the other. Second, *Ce*CP acts independently of cell cycle control to shift this balance towards mini-comet assembly.

### Capping protein suppresses assembly of reconstituted filopodia *in vitro*

A simple prediction from these *Ce*CP depletion results is that increasing CP concentrations during mitosis should shift the balance of assembly towards decreased filopodia and increased mini-comets. Our efforts to overexpress CAP-1/CAP-2 in the zygote to test this prediction were unsuccessful. Therefore, we turned to *in vitro* reconstitution methods to test how changing CP concentrations affects filopodium assembly from a branched actin precursor (Suarez et al. 2023). We reconstituted assembly of filopodia *in vitro* using micron-sized beads coated with the Arp2/3 complex activator *Ce*WVE-1 (WVE-1 pWA), and a suite of purified proteins including actin, Arp2/3 complex, profilin, plastin and formin CYK-1 (Fig. 4 A). WVE-1 pWA activates Arp2/3 complex at the bead surface to assemble branched actin filaments, some of whose barbed ends are subsequently bound and elongated by formin CYK-1. These longer filaments are bundled by plastin, creating bundles of long, formin-mediated actin filaments protruding from an Arp2/3 complex-mediated base, which we refer to as filopodia-like networks (FLNs).

**Figure 4.**
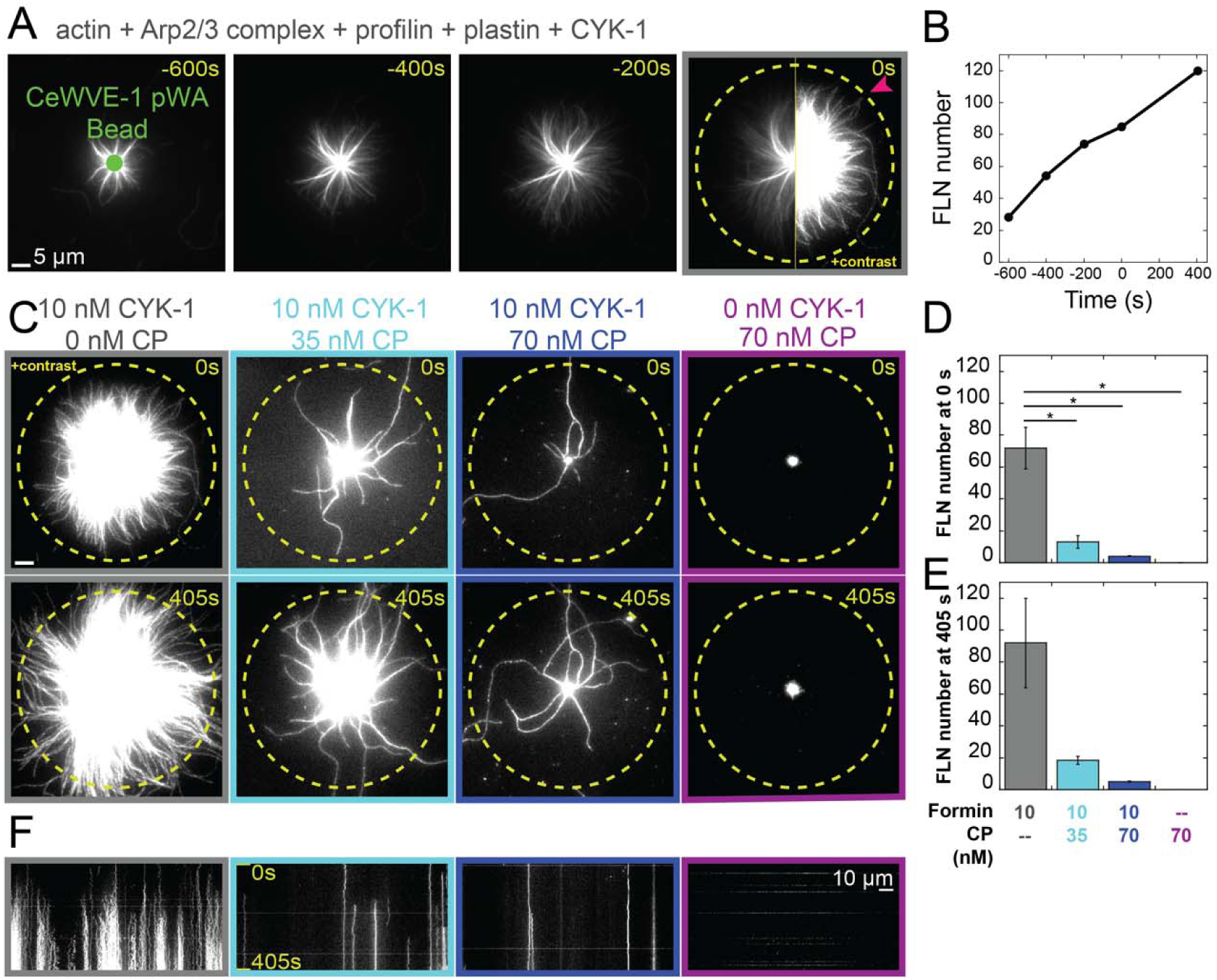
Increasing *Ce*CP reduces filopodia-like network (FLN) assembly *in vitro*. *In vitro* TIRFM visualization of the assembly of 1.5 µM Mg-ATP actin (10% Alexa488-labeled) on polystyrene beads coated with the Arp2/3 complex activator *Ce*WVE-1 pWA, in the presence of 50 nM mArp2/3 complex, 1.5 µM *Ce*PFN-1, 100 nM plastin *Ce*PLST-1, 10 nM formin *Ce*CYK-1 and a range of *Ce*CP concentrations. **(A)** Representative time-lapse images (lower image contrast) of branched actin assembly from a WVE-1 pWA-coated bead over time in the absence of *Ce*CP. t=0s is the time at which the first F-actin bundle (FLN) crosses a threshold radius 26 µm from the center of the bead (pink arrowhead, dashed yellow circle). **(B)** Number of FLNs emanating from the bead surface over time for the experiment in the absence of *Ce*CP shown in panel A. **(C)** Representative images (higher image contrast) of reactions with the indicated concentrations of *Ce*CYK-1 and *Ce*CP when the first FLN crosses the 26 µm radius (upper, 0s) and 405s later (lower). In the absence of *Ce*CYK-1 (far right), t=0s corresponds to 405s before the end of the experiment since no FLN crosses the 26 µm radius. **(D-E)** Number of FLNs emanating from the bead surface at **(D)** t=0s and **(E)** t=405s. N=2 beads for each condition. **(F)** Kymographs tracing the circumference of the 26 µm threshold from time 0 s to 405 s (top to bottom) from panel C.

We measured the number of FLNs over time (Fig. 4 A-B), at the timepoint (time 0) at which the first FLN crosses a specific length threshold (Fig. 4 A, yellow dashed circle), and 405 s after time 0. To test the effect of *Ce*CP on FLN assembly, we performed the assembly reaction in the presence of different concentrations of purified *Ce*CP (Fig. 4 C-F). In the absence of *Ce*CP, many formin-mediated FLNs emanate from the beads (Fig. 4 C, gray outlines), and the number of FLNs increases over time from 0 to 405 s (Fig. 4 D-E, gray). In the presence of 35 nM *Ce*CP the number of FLNs observed at 0 s and 405 s decreases 5-fold relative to 0 nM *Ce*CP (Fig. 4 C-E, cyan). Addition of 70 nM *Ce*CP further suppresses FLN assembly more than 10-fold (Fig. 4 C-E, dark blue). In control reactions without formin, FLNs cannot form in the presence of *Ce*CP (Fig 4 C-E, purple). Kymographs of intensity at the length thresholds (yellow dashed circles in Fig. 4C) illustrate that increasing CP decreases FLN formation, and that FLNs cannot form in the absence of formin (Fig. 4 F). These results indicate that competition between formin CYK-1 and *Ce*CP for F-actin barbed ends regulates filopodium assembly (Suarez et al. 2023).

### Capping protein depletion increases filopodium lifetime and slows filopodium assembly dynamics

To gain further insight into how capping protein regulates filopodium assembly *in vivo*, we compared filopodium morphology and dynamics in control and *cap-1(RNAi)* embryos (Fig. 5). We observed three-step dynamics for wild-type filopodium assembly. Filopodia initiate and rapidly grow to a steady-state length of ∼1.5-2 µm, which they maintain as they move directionally across the cortex, and then rapidly shrink and disappear from the cortex (Fig. 5 D, yellow, green, blue shaded areas, respectively, Fig. S5 A). Upon *Ce*CP depletion, we observed a variety of changes in filopodium morphology. To compare filopodium F-actin bundle thickness, we measured the integrated fluorescence intensity of the cross-section of all filopodia at 80s after PCR in zygotes expressing LifeAct::mCherry (Fig. 5 A, solid orange line B). We normalized each measurement to the background directly neighboring each individual filopodium (Fig. 5 A, dotted line B). The integrated intensities of these cross-sections are 1.06-fold higher in *cap-1(RNAi)* cells than in control cells (Fig. 5 B), suggesting that *Ce*CP depletion has a small but statistically significant effect on the number of actin filaments in the shafts of filopodia. Additionally, we measured the lengths of filopodia (Fig. 5 A, yellow line C) throughout their lifetimes, and found that *cap-1(RNAi)* causes a slight, but not statistically significant, increase in maximum filopodium length (Fig. 5 C).

While there are only small changes in filopodium morphology, we observed significant differences in filopodium dynamics. We compared the lifetimes of filopodia, from initiation to disassembly (Fig. 5 D), in control and *cap-1(RNAi)* embryos. *Ce*CP depletion significantly increases filopodia lifetime more than two-fold compared to controls (Fig. 5 E-F), from an average of ∼40 s in controls to an average of ∼100 s in *cap-1(RNAi)* (Fig. 5 F). In fact, most filopodia assembled by *cap-1(RNAi)* cells are longer-lived than the longest-lived filopodia in control cells (Fig. 5 E), suggesting that *Ce*CP plays an important role in the maintenance and/or disassembly of filopodia.

To compare the assembly and disassembly dynamics of filopodia, we plotted the lengths of filopodia over time as they appear and disappear from the cortex (Fig. 5 G-J, Fig. S5). Given that *Ce*CP halts F-actin barbed end assembly, and that *Ce*CP depletion significantly increases filopodium lifetime, we wondered whether *Ce*CP depletion would change filopodium assembly rates. To assess filopodium assembly dynamics, we aligned all filopodium length measurements with respect to the first frame in which each filopodium appears and plotted their length over time (Fig. 5 G, Fig. S5 A-C). In contrast to our prediction, *cap-1(RNAi)* decreases the filopodium growth rate, increasing the time to half maximum length (t_halfmax_) from ∼2 s to ∼4 s (Fig. 5 H), and decreasing the slope at t_halfmax_ threefold, from 0.36 µm/s to 0.12 µm/s (Fig. S5 D). To assess filopodium disassembly dynamics, we lined up the filopodium length measurements at the last frame before each filopodium disappears from the cortex and plotted lengths over time for the last 20 s of their lifetimes (Fig. 5 I). Again, *Ce*CP depletion decreases the disassembly rates of filopodia, increasing t_halfmax_ from ∼2 s in control cells to ∼3 s in *Ce*CP-depleted cells (Fig. 5 J), and more drastically decreasing the slope at t_halfmax_ from 0.42 µm/s to 0.19 µm/s (Fig. S5 E). Thus, *Ce*CP depletion increases filopodium lifetime and slows filopodium assembly and disassembly dynamics.

### CP’s role in filopodium assembly can be partially attributed to competition between formin and CP

While it is clear that *Ce*CP has an important role in filopodium formation, it is unclear how *Ce*CP contributes to filopodium assembly and dynamics. *In vitro,* CP competes with formins and other actin elongators for binding barbed ends. However, *in vivo,* the two ABPs often have opposing roles in actin assembly (Shekhar et al. 2015; Bombardier et al. 2015; Billault-Chaumartin and Martin 2019; Kovar et al. 2005; Harris et al. 2004; Zigmond et al. 2003; Barzik et al. 2005; Bear et al. 2002; Moseley et al. 2004). We hypothesized that the *Ce*CP depletion phenotypes, including increased numbers of filopodia, decreased mini-comet numbers, increased filopodium lifetime, and increased filopodium bundle thickness, are in part due to an increase in formin binding and elongating barbed ends that would otherwise be capped. To test this possibility, we compared the effects of single and double depletion of formin CYK-1 and *Ce*CP (Fig. 6 A-G). To control for dilution effects due to mixing of RNAi cultures in double depletion experiments, we diluted our single depletion cultures 1:1 with empty vector control RNAi culture.

**Figure 5.**
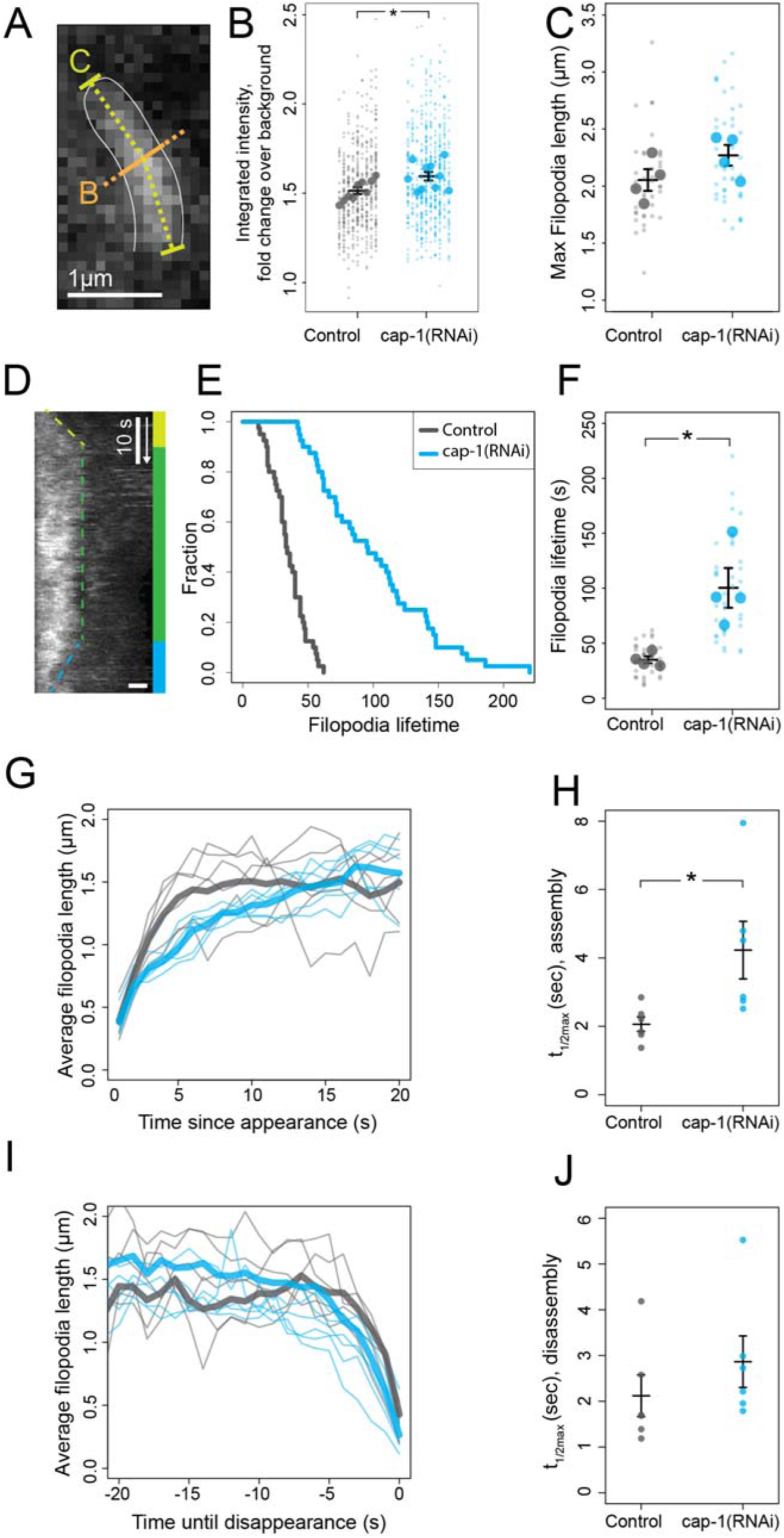
*Ce*CP depletion affects morphological and dynamic properties of filopodia. Quantification of the effects on filopodia of RNAi depletion of *cap-1* (*Ce*CP) in *C. elegans* zygotes expressing Lifeact::mCherry, visualized by near-TIRFM. **(A-C)** Morphological properties of filopodia. **(A)** Representative filopodium from a zygote expressing Lifeact::mCherry, showing measurements of F-actin thickness (orange line B) and length (yellow line C). **(B)** An estimate of the actin bundle thickness in the filopodia shaft. Integrated Intensities of F-actin signal (Lifeact::mCherry) across perpendicular cross sections of filopodia, reported as fold change over local background. Large dots represent the average of each embryo (N=9 control, N=10 *cap-1(RNAi)*). Small dots are measurements of single filopodia within an embryo. Black bars are the mean of the means ± S.E.M. p=0.018, unpaired t-test **(C)** Maximum filopodium length over their lifetimes. Large dots are the average for each embryo (N=4 for each condition), small dots are measurements of individual filopodia within an embryo (n=10 filopodia per embryo). Black bars are mean of means +/− S.E.M. p=0.15 (n.s.), unpaired t-test **(D)** Kymograph of a single representative control LifeAct::mCherry-labeled filopodium over its measurable lifetime. Time zero, at the top, is the first frame in which the filopodium appears. Yellow indicates the initiation/growth phase, green indicates the steady state/treadmilling phase, blue indicates the disassembly/shrinking phase. **(E)** Distribution of filopodium lifetimes. t=0 is the first frame of appearance of each filopodium. N=4 embryos, n=10 filopodia per embryo. **(F)** Filopodium lifetimes. Large dots are the average for each embryo (N=4 embryos for each condition), small dots are measurements of individual filopodia within an embryo (n=10 filopodia per embryo). Black bars are mean of means +/− S.E.M. p=0.034, unpaired t-test **(G)** Filopodium assembly dynamics. First 20 seconds of filopodium assembly, aligned so that t=0 is the first frame in which the filopodium appears. Thin lines are the average of 10 filopodia in a single embryo, thick lines are the averages over all embryos (N=6 embryos for each condition). **(H)** Average time it takes for filopodia to reach half their maximum length, based on nonlinear fits to the per-embryo average curves in G) (N=6 embryos, 10 filopodia per embryo). Mean of means +/− S.E.M., p=0.049, unpaired t-test **(I)** Filopodium disassembly dynamics: last 20 seconds of filopodia lifetimes. Same dataset as in E, but all measurements are aligned so that time 0 is the last frame the filopodia is visible. Thin lines represent the average of 10 filopodia in a single embryo, thick lines are the averages of all embryos (N=6 embryos for each condition). **(J)** Average time between filopodia reaching half maximum length and disappearance from the cortex. Based on nonlinear fits to the per-embryo average curves in (G) and (I)(N=6 embryos, 10 filopodia per embryo), aligned at disassembly as in H. Mean of means +/− S.E.M., p=0.33 (n.s.), unpaired t-test

**Figure 6.**
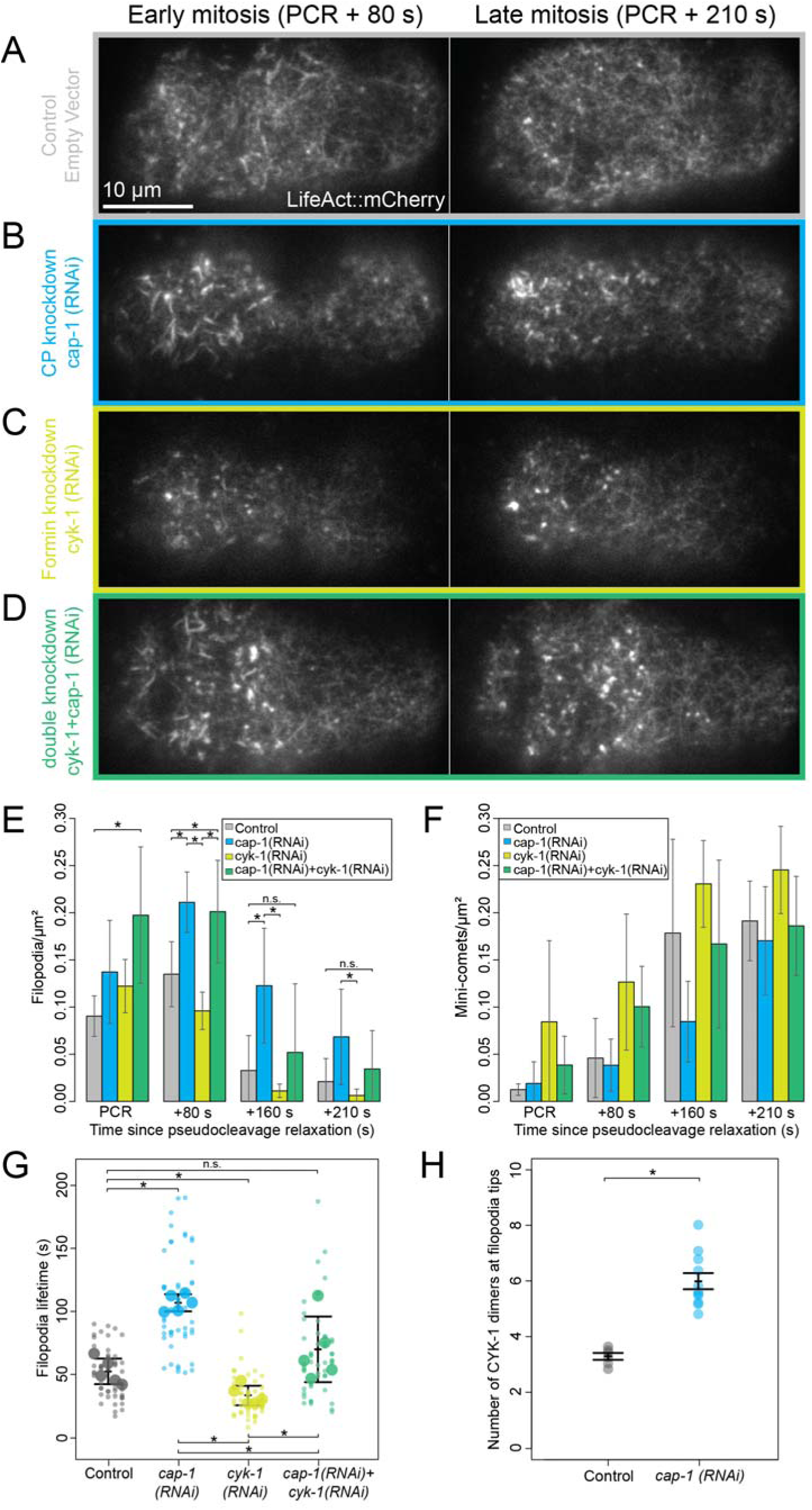
*Ce*CP regulates filopodia assembly via competition with formin CYK-1. (**A-D**) Near-TIRF images of *C. elegans* zygotes expressing LifeAct::mCherry at two timepoints: early mitosis (80 seconds after pseudocleavage furrow relaxation (PCR)), and late mitosis (210 seconds after PCR). Vertical axis labels indicate RNAi conditions. **(E)** Anterior filopodia density over time for each RNAi condition. Error bars represent standard deviation. N=5 embryos for each condition. *p<0.05, n.s.=not significant. **(F)** Anterior mini-comet density over time for each RNAi condition, graphed as in E. **(G)** Lifetimes of filopodia. Large dots represent the average of 10 filopodia per embryo, small dots represent each of the 10 filopodia measured per embryo. Error bars: mean of means, +/− S.E.M. **(H)** Estimate of the amount of CYK-1::GFP at the tips of filopodia in embryos endogenously expressing CYK-1::GFP; CAP-1::mKate; ARX-7::HALO. N=6 control embryos, N=11 *cap-1(RNAi)* embryos. Error bars: mean of means, +/− S.E.M.

We confirmed (Fig. 3) that relative to controls, partial *Ce*CP depletion increases filopodium density by 1.5 to 3.7-fold throughout mitosis and decreases mini-comet density 2.0-fold in late mitosis (Fig. 6 B, E, F, blue). Conversely, partial depletion of formin CYK-1 alone has the opposite effect relative to controls (Fig. 6 C, E, F, yellow), decreasing filopodium density in late mitosis by 3.0-fold, and increasing mini-comet density 1.3-to 6.0-fold throughout mitosis. Given that formin is not present in mini-comets (Fig. 1 E), the effect of formin depletion on mini-comet assembly is likey to be indirect.

Next, we compared the simultaneous reduction of formin CYK-1 and *Ce*CP to the depletion of formin or *Ce*CP individually (Fig. 6 A-G, green). While *Ce*CP depletion alone decreases mini-comet density compared to controls, double depletion of formin and *Ce*CP restores mini-comet density to near control levels. Interestingly, co-depletion of formin and *Ce*CP does not suppress the increase in filopodium density during early mitosis that we observe upon *Ce*CP depletion alone (Fig. 6 E, PCR, +80s). However, in late mitosis, co-depletion of formin and *Ce*CP decreases filopodium density by ∼2.0-fold compared to *Ce*CP depletion alone. The results in late mitosis are consistent with the hypothesis that competition between *Ce*CP and formin regulates filopodium assembly, as co*-*depletion of formin and *Ce*CP restores filopodium density to similar levels as control embryos (Fig. 6 E, +160 s, +210 s, Video 10). Thus, while competition between *Ce*CP and formin regulates ectopic filopodium assembly during late mitosis upon *Ce*CP depletion, it does not explain the increase in filopodium assembly in early mitosis, suggesting that different mechanisms may govern filopodia initiation in early and late mitosis.

To test whether increased filopodium thickness upon *Ce*CP depletion (Fig. 5 B) is due to an increase in formin-mediated filaments in the shaft, we measured filopodium bundle thickness upon single and double *Ce*CP and formin depletions Fig. S6 A). Because the RNAi is diluted in double RNAi experiments, depletions are not strong enough to reproduce the significant increase in filopodium bundle thickness (Fig. 5 B). However, there is a small increase in bundle thickness upon *Ce*CP depletion, and a small decrease upon formin depletion during early mitosis (PCR+80s) (Fig. S6 A), bolstering that formin and CP have opposing roles in determining the thickness of filopodia. As a more direct test of increased formin activity in filopodia upon *Ce*CP depletion, we compared the relative amounts of formins at the tips of filopodia in control and *cap-1(RNAi)-*treated embryos (Fig. 6 H). We imaged filopodia tips in cells with endogenously-labeled formin, then photobleached them to single-molecule level. Using the fluorescence intensity of a single molecule as a reference, we estimated the number of formin dimers at the tips of filopodia and found that *Ce*CP depletion significantly increases the amount of formins at the tips of filopodia by ∼2.0-fold (Fig. 6 H). Consistent with our hypothesis that formin and *Ce*CP are in competition with each other, this result supports a model in which formin activity increases upon *Ce*CP depletion.

We also tested whether competition between *Ce*CP and formin tunes filopodium lifetime. We measured the lifetimes of filopodia in control, *Ce*CP RNAi, formin RNAi, and *Ce*CP/formin double RNAi conditions. As before, *Ce*CP depletion increases filopodium lifetime, whereas formin depletion decreases filopodium lifetime (Fig. 6 G). Depletion of both *Ce*CP and formin rescues filopodium lifetimes to control-type levels (Fig. 6 G), indicating that the proportional amounts of these two proteins is a better determinant of filopodium lifetimes than the total amounts of either CP or formin. This result suggests that competition between formin and CP for binding F-actin barbed ends regulates the lifetimes of filopodia.

One possible mechanism by which *Ce*CP could help limit filopodium lifetime is by triggering filopodium disassembly, possibly by competing formin off the elongating F-actin barbed ends at the tips of filopodia. To test whether *Ce*CP displaces formin at filopodium tips to trigger filopodium disassembly, we made line scans along the lengths of filopodia in zygotes expressing ARX-7::HALO (Arp2/3 complex), CYK-1::GFP (Formin); CAP-1::mKate (*Ce*CP), and measured the fluorescence intensity at the tips of the filopodia over time (Fig. S6 B). While CYK-1 intensity decreases in the last 10 seconds of filopodium lifetime, we do not detect an increase in CP localization near filopodium tips preceding filopodia disassembly (Fig. S6 B).

An alternative hypothesis is that *Ce*CP and formin compete at the filopodium base to regulate the elongation of filaments into the shafts of filopodia and thus the maintenance of the structure. We explored this possibility by imaging filopodia in zygotes expressing endogenously tagged formin CYK-1::GFP, and LifeAct::mCherry at very high frame rates (10 fps) (Fig. S6 C, Video 11). In addition to bright puncta at the tips of filopodia, we observed highly dynamic formin punctae along their shafts (Video 12). From kymographs of these filopodia over time, we observed rapid directional movements of formin punctae towards the tips of filopodia throughout their lifetimes. (Fig. S6 C, yellow arrows). Importantly, these punctae move at speeds similar to those observed for formin dimers elongating individual actin filaments within the cortical meshwork (Li and Munro 2021). Therefore, we hypothesize that these formins assemble new actin filaments to add to the filopodia bundle, and that these events may be important for maintaining filopodium assembly. Thus, competition between formin and *Ce*CP for binding filaments at the bases of filopodia could regulate the frequency of these elongation events and may contribute to changes in filopodium lifetimes (Fig. 7 C). However, measurements of the frequency of formin movements towards the tips of filopodia do not reveal a significant difference between control and *Ce*CP-depleted filopodia (Fig. S6 D).

**Figure 7.**
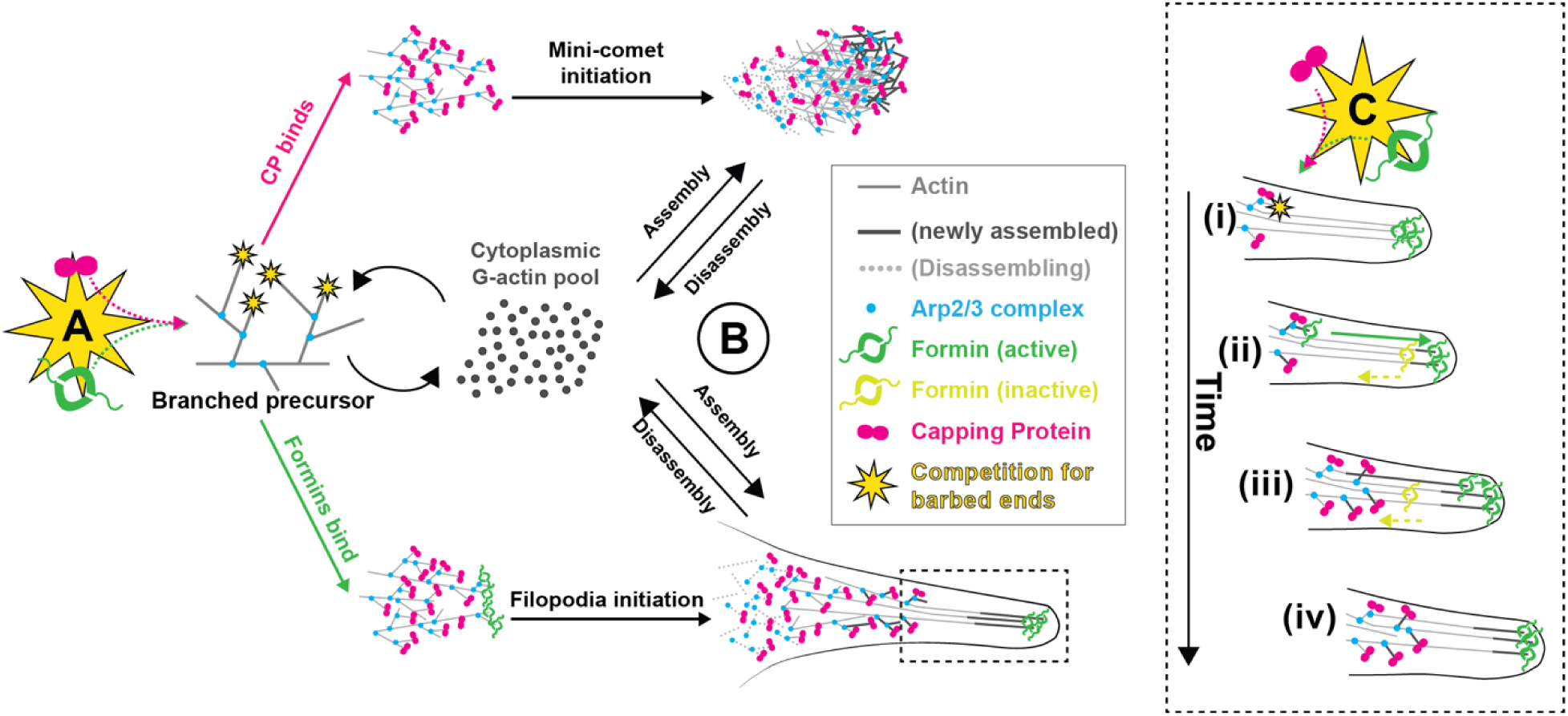
Competition between *Ce*CP and formin for barbed ends plays multiple roles in internetwork competition and filopodium maintenance. (**A**) Initiation: *Ce*CP and formin CYK-1 compete for binding the free barbed ends of Arp2/3 complex-assembled branched precursor networks. If *Ce*CP activity dominates (pink arrow), increased branching leads to mini-comet assembly. If formin activity is high (green arrow), barbed ends are elongated and bundled to form the protrusive shaft of a filopodium. **(B)** Filopodia and mini-comets are treadmilling structures that undergo continuous assembly and disassembly through a common cytoplasmic pool of actin monomers. **(C)** Competition between formin and *Ce*CP regulates filopodia maintenance. (i) *Ce*CP and formin compete to bind a new barbed end at the base of a filopodium. (ii) Formin associates and the filament rapidly elongates (dark green), whereas another formin is stochastically inactivated at the tip of the filopodium (light green). (iii) Elongation of active formin-associated barbed ends (dark green) continue to extend the filopodium and the inactive formin-associated filament (light green) is left behind. (iv) Filopodium assembly and extension persists.

## Discussion

It is well-established that capping protein (CP) regulates actin filament length by associating with and inhibiting barbed end dynamics (Isenberg et al. 1980; Wanger and Wegner 1985; Caldwell et al. 1989; Cooper et al. 1984; Cooper and Pollard 1985). However, the mechanisms by which filament length control generates distinct F-actin network architectures, and helps to ensure the proper allocation of limiting resources like G-actin, remain poorly understood. Here, we uncovered key roles for *C. elegans* capping protein (*Ce*CP) in balancing F-actin assembly, both within and across different F-actin networks in the *C. elegans* zygote. We focused on two prominent F-actin networks, filopodia and mini-comets. We provided new detailed characterization of their components, dynamics, and mechanisms of assembly. We found that *Ce*CP helps regulate the relative abundance of these networks and propose that competition between *Ce*CP and the actin assembly factor formin within a common branched precursor dictates network architecture, generating either a filipodium or a mini-comet (Fig. 7 A). In addition to balancing the assembly of filopodia and mini-comets, we found that competition between formin and *Ce*CP for barbed ends also regulates filopodia dynamics and lifetime (Fig 7 C). Additionally, our data suggest that filopodia and mini-comets compete for a shared pool of resources, such as G-actin. Therefore, competition between *Ce*CP and formin allocates limited resources toward either Arp2/3 complex– or formin-mediated assembly, expanding filament– and network-level competition into global F-actin cytoskeleton regulation (Fig. 7 B). This study establishes that *Ce*CP is an important regulator of F-actin network fate, F-actin network architecture and dynamics, and a determinant of global G-actin allocation among different cortical F-actin networks.

### *Ce*CP is enriched in filopodia and mini-comets

Live imaging of endogenously labeled proteins revealed that *Ce*CP is a component of the major cortical F-actin networks in the zygote. *Ce*CP is highly enriched in the Arp2/3 complex-assembled domains of filopodia and mini-comets, but is more weakly associated with the formin-mediated cortical network and contractile ring (Fig. 1 B-D). Filopodia were previously described in *C. elegans* zygotes as membrane-bound protrusions consisting of plastin-bundled F-actin shafts with formins at their tips (Hirani et al. 2019). We found that like canonical filopodia found in many other systems (Mattila and Lappalainen 2008), *C. elegans* filopodia have Arp2/3 complex-assembled bases that are dense, motile clouds of branched F-actin, enriched in *Ce*CP (Fig. 1 C). Filopodium bases closely resemble the Arp2/3 complex– and *Ce*CP-rich mini-comets, suggesting that they share a common precursor (Fig. 7 A). Filopodium bases and mini-comets both exhibit treadmilling motility, as observed at the leading edges of many motile cells (Blanchoin et al. 2014). Filopodia and mini-comets move with similar speeds, although filopodia move with greater directional persistence (Videos 1-2). Therefore, Arp2/3 complex-mediated branched F-actin assembly appears to be the common engine driving the motility of both filopodia and mini-comets (T. Svitkina 2018), while formin-mediated assembly at the tips of filopodia may contribute to their directional persistence. The local coexistence of these two structures at the surface of a single large cell provides a unique opportunity to study how CP modulates the interplay between formin– and Arp2/3 complex-mediated F-actin assembly, both within a single composite structure (filopodia) and across different networks (between filopodia and mini-comets).

### Competition between *Ce*CP and formin CYK-1 regulates the balance of filopodia and mini-comet assembly

We showed that depleting *Ce*CP induces an increase in filopodia, and a decrease in mini-comets throughout the cell cycle, including in late mitosis when filopodia are normally absent and mini-comets are abundant (Fig. 3 A-B). Conversely, formin CYK-1 depletion induces fewer filopodia and more mini-comets (Fig. 6 E-F). These results suggest that *Ce*CP and formin exert opposing influences on a switch between filopodia and mini-comet assembly (Fig. 7 A), consistent with the two proteins having opposing roles in other systems (Kovar et al. 2005; Zigmond et al. 2003; Harris et al. 2004). Importantly, simultaneous depletion of *Ce*CP and formin has almost no effect on filopodium and mini-comet abundance in late mitosis, suggesting that the ratio of CP to formin activity controls the switch that determines assembly of a filopodium or a mini-comet (Fig. 6 E-F).

Our data suggest that direct competition for barbed ends by CP and formin regulates a network-level decision to initiate filopodium or mini-comet assembly in the *C. elegans* zygote (Fig. 7 A), as has been postulated in a range of other biological contexts (Shekhar et al. 2015; Bombardier et al. 2015; Harris et al. 2004; Zigmond et al. 2003; Moseley et al. 2004). We propose a mechanism for filopodia and mini-comet assembly that starts with a common precursor network: Arp2/3 complex-mediated branched filament structures with free barbed ends that begin to appear in interphase (Fig. 7A). We propose that these precursor networks are akin to the branched F-actin networks from which filopodia emerge in other systems through convergent elongation (T. M. Svitkina et al. 2003; Mejillano et al. 2004) and form either filopodia or mini-comets based on the balance of *Ce*CP and formin activity at barbed ends. *Ce*CP-dominated activity promotes dense branching (Akin and Mullins 2008) leading to mini-comet assembly. Conversely, formin-dominated activity promotes elongation of linear filaments, which are bundled by plastin to create the shaft of a filopodium (Fig. 7 A). In this way, the relative activities of *Ce*CP and formin regulate a switch that determines whether a filopodium or a mini-comet is assembled from branched precursor networks.

Competition between CeCP and formin facilities the proper balance of filopodia and mini-comets in late mitosis. However, given that double depletion of CeCP and formin does not restore wild-type levels of F-actin networks in early mitosis (Fig. 6 E and F, PCR and PCR +80), additional regulatory mechanisms likely exist. Given that *Ce*CP associates with barbed ends for the lifetime of a filament (Fig. 2 E-H), *Ce*CP activity is likely controlled through modulation of the on-rate *Ce*CP binds barbed ends. Part of this regulation is via competition with formin, but there are also multiple regulators of CP activity expressed in the *C. elegans* zygote, including V-1/Myotrophin (MTPN-1), CARMIL (CRML-1), and twinfilin (TWF-2) that may further modulate CP activity (Edwards et al. 2014; Saini et al. 2025).

### Competition between *Ce*CP and formin within filopodia regulates their size and assembly dynamics

Upon *Ce*CP depletion, in addition to an increased abundance, filopodia are longer-lived, thicker, and contain more formin at the tips (Fig. 6 H, Fig. 5 B and F). This indicates that *Ce*CP has additional roles in regulating filopodium morphology and dynamics. Given that filopodia are shorter-lived upon formin depletion and double depletion of formin and *Ce*CP restores filopodium lifetime back to control levels (Fig. 6 G), we propose that competition between *Ce*CP and formin for barbed ends helps regulate filopodia thickness and lifetime (Fig. 7 C). One possibility is that *Ce*CP competes with formins for barbed ends at filopodia tips. However, we do not observe an increase of *Ce*CP at filopodia tips prior to their disassembly (Fig. S6 B). Although this may be due to imaging limitations, we favor an alternative mechanism whereby *Ce*CP helps regulate filopodium morphology and lifetime by competing with formin for barbed ends at the branched network base (Fig. 7 C). High-speed imaging allowed direct observation of formins moving from the bases to the tips of filopodia throughout their lifetimes (Fig. S6 C). We posit that formins bind barbed ends in the branched base and elongate actin filaments along the shafts of filopodia to ensure their mechanical integrity, which maintains the stiffness to push against and deform the cell membrane. Thus, competition between *Ce*CP and formin regulates the number of elongating filaments in filopodial shafts. If formin competes sufficiently with *Ce*CP, new formin-assembled filaments will replace stochastically inactivated ones, and filopodium assembly will be maintained (Fig. 7 C). Conversely, if *Ce*CP sufficiently out-competes formin, then the number of formin-assembled filaments in the shaft will diminish over time, terminating filopodium assembly. Thus, by regulating the frequency with which formins elongate new barbed ends from the base, *Ce*CP helps regulate the thickness and the lifetimes of filopodia.

*Ce*CP depletion also slows the initial filopodium elongation rate (Fig. 5 G-H), which may be due to an increased number of elongating barbed ends in the confined, membrane-bound shaft. Upon initiation, filopodium tips elongate faster than the base until they reach a stereotypical terminal length of ∼2 µm, which then stays relatively constant until termination (Fig. 5 D and S5). The slowing of filopodium shaft assembly as filopodia approach a terminal length may reflect a threshold for membrane resistance and/or decreased actin monomer availability with increasing length, consistent with previous models of filopodium length regulation (Mogilner and Rubinstein 2005; Lan and Papoian 2008).

### *Ce*CP indirectly regulates global F-actin cytoskeleton assembly by controlling resource allocation among different modes of assembly

Competition between *Ce*CP and formin may also indirectly affect the assembly of other F-actin networks (Suarez and Kovar 2016). In other systems the amounts of actin monomers, ABPs, or upstream actin regulators within a cell are limiting, and competition between F-actin networks for these resources helps regulate their size and density (Suarez and Kovar 2016; Kadzik et al. 2020). Therefore, *Ce*CP’s role in regulating filopodium abundance and size could help balance the allocation of G-actin and/or other resources to competing F-actin networks in the zygote (Fig. 7 B). The abundance of filopodia and mini-comets are reciprocally affected upon depleting *Ce*CP or formin, whereby over-assembly of one leads to under-assembly of the other. While a limited number of branched precursors could explain changes in filopodium and mini-comet abundance, we suspect that competition for limited resources is the primary driver. Over-or under-assembly of filopodia either depletes or frees up resources available for mini-comet assembly. Consistent with this model, although formin is not associated with mini comets (Fig. 1 E iii-iv), formin depletion increases mini-comet assembly in late mitosis (Fig. 6 F). Thus, our data suggest that the cortical F-actin networks in the *C. elegans* zygote are in homeostatic balance for actin monomers and/or ABPs, and competition between *Ce*CP and formin robustly modulates this balance by biasing resource allocation to either filopodia or mini-comets. (Fig. 7 B). While we focused on filopodia and mini-comets, we predict that a similar competition for resources may govern the relative abundances of other coexisting F-actin networks in the zygote and later development stages (Chan et al. 2019). In this study, we show that through its ability to compete with formins for individual barbed ends, CP directly shapes the architecture and assembly dynamics of individual networks, as well acting as a gatekeeper for the allocation of resources among multiple networks. Thus, through its ability to control the fates of single barbed ends, CP, and other determinants of barbed end dynamics, may indirectly regulate all coexisting F-actin networks in a cell.

## Materials and methods

### C. elegans culture

We cultured *C. elegans* strains under standard conditions (Brenner 1974) on 60 mm petri plates containing E coli bacteria raised on normal, growth medium. Table 1 lists the strains used in this study. Unless otherwise specified, strains were provided by the Caenorhabditis Genetics Center, which is funded by the National Center for Research Resources.

### RNA interference

We performed RNAi using the feeding method (Timmons et al. 2001). We obtained the feeding strain targeting GFP from Jeremy Nance, the feeding strain targeting CAP-1 in the T444T feeding vector from Ronen Zaidel-Bar, and all other feeding strains from the Ahringer library (Kamath et al. 2003). To prepare RNAi plates, we grew bacteria to log phase in LB with 50µg/ml ampicillin, seeded ∼200µl of culture onto NGM plates supplemented with 100µg/ml ampicillin and 1mM IPTG, set the plates at room temperature for two days, and then stored them at 4°C for up to one week before use.

For all feeding experiments, we transferred L4 larvae onto seeded RNAi plates and cultured them at 20°C for the following amounts of time: *cap-1(L4440)(RNAi)* (46-48 h), *cap-1(T444T)(RNAi)* (25-30 h)*, cyk-1(RNAi)* (41-46 h), *gfp(RNAi)* (21-22h). To achieve the strongest *cap-1(RNAi)* phenotypes possible for both single and double RNAi experiments, we determined the longest time on RNAi plates that still allowed for production of zygotes. We confirmed moderate depletion of CYK-1 by observing cortical instabilities during interphase. We used bacteria expressing the appropriate empty vectors (L4440 or T444T) as paired controls for each experiment; we prepared control and experimental feeding plates under the same conditions and compared embryos from worms cultured on control and experimental feeding plates for the same range of times.

### Protein expression and purification

SNAP-*Mm*CP (*Mm*CapZα1-6xHIS; SNAP-CapZβ2) vector (pRSFDuet-1, pBJ 2041) was a gift from the Cooper lab (Kim et al. 2012). SNAP-*Ce*CP (*Ce*CAP-1-6xHis; SNAP-*Ce*CAP-2) vector was made by cutting out CapZα1 from the SNAP-*Mm*CP vector via restriction digest and inserting *Ce*CAP-1 gBlock (IDT) (sequence from wormbase.org, codon optimized using IDT Codon Optimization tool) via gibson assembly (NEB). The whole vector except CapZβ2 was then amplified by PCR, and *Ce*CAP-2 was inserted via gibson assembly. The final plasmid sequence was confirmed by sequencing through Plasmidsaurus.

Recombinant *C. elegans* (dual expression construct of *Ce*CAP-1-6xHis and SNAP-*Ce*CAP-2; SNAP-*Ce*CP) and mouse CP (dual-expression construct of *Mm*CapZα1-6xHIS and SNAP-CapZβ2; SNAP-*Mm*CP) proteins were purified from *Escherichia coli* strain BL21-Codon Plus (DE3)-RP (Agilent Technologies), after expression with 0.5 mM isopropyl β-d-1-thio-galactopyranoside for 16 h at 16°C. Cells were lysed by sonication in CP extraction buffer (50 mM NaH_2_ PO_4_, pH 8.0, 500 mM NaCl, 10% glycerol, 10 mM imidazole, 10 mM β-mercaptoethanol, and 1 cOmplete, EDTA-free Protease Inhibitor Cocktail tablet per 50 mL (Roche)), and clarified by centrifugation at 16,000 rpm for 15 min, followed by 18,000 rpm for 30 min, at 4° C in a Beckman SS-34 rotor. The extract was incubated for 1 hour at 4°C with Talon Resin (Takara Bio Inc.), loaded onto a column, washed with extraction buffer, and eluted with 250 mM imidazole. SNAP-CP eluent was dialyzed against CP buffer (10 mM Tris, pH 7.5, 40 mM KCl, 50% glycerol, 0.5 mM DTT, and 0.01% NaN_3_). SNAP-CP was labeled with SNAP Surface 647 according to the manufacturers’ protocols (New England Biolabs, Inc.). Concentrations of SNAP-tagged proteins and the degree of labeling were determined by measuring absorbance at 280 nm using ε_280_ =94770 M/cm (SNAP-*Ce*CP) and ε=97270 M/cm(SNAP-*mm*CP) and fluorophore absorbance using the extinction coefficient of SNAP-surface647: ε_650_ =250,000 M/cm. CP was flash-frozen in liquid nitrogen and stored at −80°C. *Gg*Actin was purified from chicken skeletal muscle acetone powder by a cycle of polymerization and depolymerization and gel filtration (Spudich and Watt, 1971). Gel-filtered actin was labeled on surface lysines with Alexa Fluor 488 succinimidylester (Thermo Fisher Scientific; (McCullough et al. 2011; Kang et al. 2012)).

*Ce*Actin was purified from N2 *C. elegans* pellets as described in (Ono and Pruyne 2012). Briefly, frozen worm pellets were thawed in homogenizing buffer (50 mM NaCl, 1 mM EDTA, 1 mM DTT, 1 mM PMSF, 20 mM Tris–HCl, pH 8.0), lysed in an Emulsi-Flex-C3 (Avestin), and centrifuged at 10,000xg for 10 minutes. Pellets were washed in homogenizing buffer and resuspended in extraction buffer (0.6 M KCl, 5 mM ATP, 5 mM MgCl_2_, 0.2 mM EGTA, 1 mM DTT, 1 mM PMSF, 20 mM Tris–HCl, pH 8.0). Extract supernatant (with 0.5% Triton X-100) was ultracentrifuged at 100,000xg for 2h. The pellet was collected and homogenized in a Dounce with 2 M Tris–HCl, pH 7.0, 1 mM MgCl_2_, 0.2 mM ATP, 0.2 mM DTT, and clarified at 100,000xg for 1 hour. The supernatant was applied to a sephacryl-300 HR column, equilibrated and run in 1 M Tris–HCl, pH 7.0, 0.5 mM MgCl_2_, 0.2 mM ATP, 0.2 mM DTT. Actin-containing fractions were pooled and dialyzed against Ca^2+^-Buffer G (2 mM Tris–HCl, pH 8.0, 0.2 mM CaCl_2_, 0.2 mM ATP, 0.2 mM DTT) overnight. Actin was then polymerized with 30 mM KCl, 2 mM MgCl_2_ and 1 mM ATP for 4 h at room temperature or overnight at 4 °C. F-actin was pelleted by ultracentrifugation at 100,000xg for 2 hours. The pellet was solubilized in G-buffer, and actin was depolymerized by dialysis against G-buffer overnight and ultracentrifuged at 424,000xg for 1 hour to remove oligomers.

CePFN-1, CePLST-1, SNAP-CYK-1(FH1FH2), and WVE-1 pWA were expressed in *E. coli* BL21-Codon Plus (DE3-RP) cells. CePFN-1 was purified via poly-L-proline affinity chromatography (Lu and Pollard 2001). CePLST-1 and SNAP-CYK-1(FH1FH2) were purified using Talon metal affinity and anion exchange chromatography (Neidt et al. 2008; Skau and Kovar 2010). WVE-1 pWA was purified by glutathione-agarose affinity chromatography (Machesky et al. 1999). Arp2/3 complex was isolated from calf thymus (Egile et al. 1999).

### Preparing glass slides and coverslips for *in vitro* TIRFM

Microscope slides (Fisher Scientific) and coverslips (#1.5, Fisher Scientific) were washed for 7 min in acetone, isopropanol, and water, followed by sonication in isopropanol for 30 min. Washed glass was incubated with piranha solution (66.6% H_2_SO_4_, 33.3% H_2_0_2_) for 2 hours, then washed with diH_2_0 and dried with air. Immediately following piranha cleaning, glass was passivated by incubation with 1 mg/mL PEG-Si (5000 MW) in 95% ethanol for 18 hours (Winkelman et al. 2014). After glass was rinsed with ethanol and water, flow chambers were prepared using double-sided tape between the slide and the coverslip.

### *In vitro* TIRFM image acquisition

For *in vitro* TIRFM experiments, we acquired images on an Olympus IX-71 microscope with an iXon EMCCD camera (Andor Technology) and a cellTIRF 4Line system (Olympus), using through-the-objective TIRF illumination. 1.5 µM 10% Alexa 488-labeled *C. elegans* Mg-ATP-actin was mixed with purified SNAP-647-CP and polymerization buffer (10 mM imidazole, pH 7.0, 50 mM KCl, 1 mM MgCl_2_, 1 mM EGTA, 50 mM DTT, 0.2 mM ATP, 50 µM CaCl_2_, 15 mM glucose, 20 µg/mL <u>catalase</u>, 100 µg/mL <u>glucose oxidase</u>, and 0.5% 400 centipoise methylcellulose) to induce F-actin assembly. Images were captured every 5 seconds using 50 ms exposure with 488 nm and 640 nm lasers.

### Bead Assay

Carboxylated 2 µm latex beads (2.6% solids; Polysciences, Eppelheim, Germany) were coated with 10 µM WVE-1 pWA (Loisel et al. 1999) and added to actin polymerization reactions containing proteins of interest, as specified in figure legends (Winkelman et al. 2016). Experiments were imaged by total internal reflection fluorescence microscopy (TIRFM).

### Pyrene assays

Pyrene seeded assembly assays were performed as previously described (Zimmermann et al. 2016). Briefly, 400 nM unlabeled preassembled actin filaments were incubated with 0.5 µM 10% *N*-(1-pyrenyl)iodoacetamide (pyrene) labeled actin monomers (labeled at Cys374) under polymerizing conditions:1x KMEI (50 mM KCl, 1 mM MgCl_2_, 1 mM EGTA, 10 mM imidazole, pH 7.0), 1x ME (50 µM MgCl_2_, 1 mM EGTA, pH 7.0), in the presence of varying concentrations of CP. Pyrene disassembly assays were performed as previously described (Zimmermann et al. 2016). Briefly, 5 µM 50% pyrene-labeled F-actin seeds were diluted to 0.1 µM in 1x KMEI and 1x ME buffers, in the presence of varying concentrations of unlabeled SNAP-CP. For both seeded assembly and disassembly assays, fluorescence (excitation= 365 nm, emission=407 nm) was measured every 10 seconds using a Tecan Infinite M200 Pro plate reader (Tecan).

### C. elegans imaging

To perform near-TIRFM imaging of embryos expressing only LifeAct::mCherry or CAP-1::GFP (for single molecule analysis), we used an Olympus IX50 inverted microscope equipped with an OMAC two-color TIRF illumination system housing 50 mW 488 and 561 nm solid-state Sapphire lasers (Coherent), and a CRISP autofocus module (Applied Scientific Instrumentation). We acquired images using a 1.45 NA oil immersion 100x TIRF objective, with 1.6X additional magnification, onto an Andor iXon3 897 EMCCD camera, yielding a pixel size of 100 nm. We set laser illumination angle manually during each imaging session to approximately maximize signal-to-noise ratio. Images were collected using Andor IQ software. For all other near-TIRFM imaging, we used a Nikon Ti2-E inverted microscope equipped with perfect focus and a LunF XL laser combiner with solid state 488 nm, 561 nm, and 640 nm lasers feeding 3 separate TIRF illumination arms. Images were acquired with a 100x 1.49 NA oil-immersion TIRF objective and 1.5X additional magnification onto an Andor IXON-L-897 EM-CCD camera yielding a pixel size of 106 nm. Laser power, illumination direction and angle, and image acquisition were controlled by Nikon Elements software. For each strain, we determined laser illumination angles empirically to maximize signal-to-noise ratio, and then used the same laser angles for all subsequent experiments. Embryos were oriented with the anterior-posterior axis perpendicular to the axis of TIRF illumination to avoid the confounding effects of small intensity gradients produced by near-TIRF illumination.

Additional image acquisition details (laser power, exposure times and frame rates) for individual experiments are provided below and in the figure legends.

### Image analysis

All image analysis was performed using FIJI (https://fiji.sc/). Unless otherwise specified below, all image analysis was performed on raw image data, and Images were processed only for displaying in the figures; none were processed before analysis.

### Estimating cortical residence times of single CAP-1::GFP molecules

To estimate cortical residence times of single CAP-1::GFP molecules, we treated worms expressing transgenic CAP-1::GFP with *gfp(RNAi)* for 21-22 hours to reduce maternal expression of CAP-1::GFP to levels where single-molecules could be detected. For each embryo, we fixed the unit exposure (100% laser power; 50 ms exposure time),and acquired images during mitosis at either low duty ratio (dr_low_= 0.1; 2 exposures/s) for 150 seconds, or at high duty ratio (dr_hi_= 1; continuous exposure) for 50 seconds.We performed single molecule tracking as described (Robin et al. 2014), using a Matlab implementation (http://people.umass.edu/kilfoil/downloads.html) of the Crocker-Grier method (Crocker and Grier 1996) to detect single molecules as local peaks in fluorescence intensity, and then µTrack software (http://github.com/DanuserLab/u-track) to track single molecules over time (Jaqaman et al. 2008). We pooled all trajectories collected from multiple embryos at low duty ratio or high duty ratio for further analysis. From these pooled data, we constructed single molecule release curves by plotting the number of tracked molecules with lifetime > T as a function of T. We used Matlab’s non-linear least squares fitting function ***nlinfit*** to fit these release curves to a weighted sum of two exponential functions:

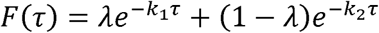

on the interval 2 s < r< ∞, and took the slower of the two rate constants as an estimate of the observed disappearance rate, which we assume is the sum of rate constant for the dissociation of F-actin-bound CAP-1 from the cortex and the photobleaching rate, which depends on duty ratio:

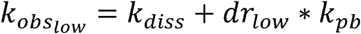

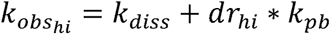

where k_pb_ is the photobleaching rate during a unit exposure. We solved the above equations simultaneously to estimate the two unknowns (k_diss_ and k_pb_). To compute confidence intervals for these estimates, we used bootstrapping to sample the pooled data 1000 times, computed estimates of k_diss_ and k_pb_ from each sample, and then took 2 times the standard deviation of the estimates as an estimate of the 95% confidence interval. All scripts used for this analysis are available upon request.

### Using pseudocleavage furrow relaxation as a time reference to compare measurements across embryos

We used pseudocleavage furrow relaxation (PCR) as a reference timepoint to compare data across embryos. We show here that pseudocleavage furrow relaxation (PCR) is a well-defined marker for cell cycle progression from interphase to mitosis: In embryos imaged in an equatorial plane, we observe a strong correlation between the timing of PCR and nuclear envelope breakdown (NEBD) with a mean delay of 231.5 s. Extending interphase by activating the DNA replication checkpoint, delays PCR and NEBD, without affecting the mean delay between them. However, unlike NEBD, which can only be observed in an equatorial image plane that contains the nuclei, the onset of PCR can also be identified in a cortical image plane, making it more suitable as a reference timepoint for experiments involving cortical imaging.

In surface (cortical) views of the zygote, the pseudocleavage furrow is visible as a local indentation of the slightly compressed cortical surface, and PCR can be visualized as the regression of these indentations. Thus, we identified the onset of PCR by identifying the first frame in which the onset of the regression can be clearly identified.

### Measuring mini-comet and filopodia numbers, thickness, length and lifetime

We performed these measurements on timelapse data acquired from embryos expressing LifeAct::mCherry during the window of time spanning pseudocleavage relaxation to cytokinesis onset, with 40% laser power, 100 ms exposure time, with 200 ms between each frame. We defined filopodia operationally as local enrichments of F-actin that: (a) exhibit processive and directional motion that can be tracked by eye over time; (b) have elongated structures for at least part of their lifetime. To count the filopodia present in a given frame, we first identified candidates (local F-actin enrichments) within that frame, and then examined the time lapse sequence around that frame to determine whether it met the above criteria. We defined mini-comets as local enrichment of F-actin that do not exhibit directionally persistent motion and stay mostly punctate throughout their lifetime. With labeled CYK-1::GFP, filopodia were only counted if they have a GFP punctum at the tip.

To measure filopodia lifetimes and lengths over time for a given embryo, we identified all filopodia that were present at 80 s after PCR. For each filopodium, we determined the frame in which it first appears and frame in which disappears and used these to compute the lifetime. For each frame over that filopodium’s lifetime, we constructed a segmented line ROI with a width of 2 pixels along the filopodium and aligned with its tip. We stacked these ROIs to create the kymograph in Fig 5 D showing intensity along the filopodium vs time. From this kymograph, we measured the filopodium length vs time, and computed the maximum length. To perform further analysis, we averaged length vs time data over ten filopodia per embryo, aligning the data across filopodia with respect to time of appearance (to measure or plot the initial growth kinetics), or the time of disappearance (to measure or plot the disappearance kinetics). To estimate the half times T_1/2_ for growth or disappearance, we used nonlinear least squares method to fit the first (or last) 20 s of the aligned length vs time data to an exponential relaxation function of the form:

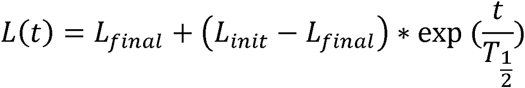

To estimate filopodia bundle thickness, we identified all filopodia in a single frame. For each, we constructed a rectangular ROI perpendicular to the filopodium with a minimum length of 9 pixels. We measured the fluorescence intensity profile along the ROIs, and identified the highest intensity pixel as the middle of the bundle. We measured the average intensity in a five-pixel window, centered on the peak. We estimated the local background fluorescence as the average intensity over two pixels flanking the measurement window on either side. Then we took our estimate of thickness as the ratio of the measured/background intensities.

To detect fast anterograde formin movements (Fig. S6), we imaged embryos expressing LifeAct::mCherry; CYK-1::GFP, with 50 ms exposure times, 10 frames per second, 3% laser power 488, 12% laser power 561. We made kymographs of representative filopodia, lined up at the tips (Fig. S6 C), and identified events as CYK-1 enrichments that appear along their shafts and move toward their tips for at least 20 frames.

## Statistical analysis

Statistical analysis was performed in Graphpad T-test calculator, unpaired or paired t-tests as indicated. P values <0.05 were reported statistically significant (*).

## Section describing supplemental material

Fig. S1 shows filopodia and mini-comet abundance and CAP-1::mKate localization to the contractile ring in control embryos. Fig. S2 shows exponential fit to single-molecule *in vivo* data and distributions of fast-diffusing and cortex-binding events. Fig. S3 shows that both control and *cap-1(RNAi)*-induced ectopic filopodia are membrane-bound, with formin CYK-1::GFP at their tips and ARX-7::HALO at their bases. Fig. S4 compares cell cycle progression in control and *cap-1(RNAi)* embryos. Fig. S5 shows full graph of filopodia lengths over time, and slopes at T_halfmax_ for filopodium assembly and disassembly. Fig. S6 A Shows filopodium widths upon CAP-1/CYK-1 double knockdown. B. shows no CAP-1::mKate enrichment at filopodia tips, Fig. S6 C-D compares frequency of anterograde formin movements in filopodia of control and *cap-1(RNAi)*-treated embryos.

## Acknowledgements

We thank members of the Kovar and Munro labs for helpful discussions, feedback, and support. This work was supported by National Institutes of Health Grants R35 GM153234 (to D.R.K.), R01 GM143576 (to E.M.) and T32 GM0071832 (to S.E.Y.), and Israel Science Foundation grant 767/20 (to R.Z.B.).

