## Supplemental Material for "Capping protein regulates the balance of assembly among diverse actin networks in *C. elegans* zygotes"

**Supplemental materials**

**Supplemental figures:**

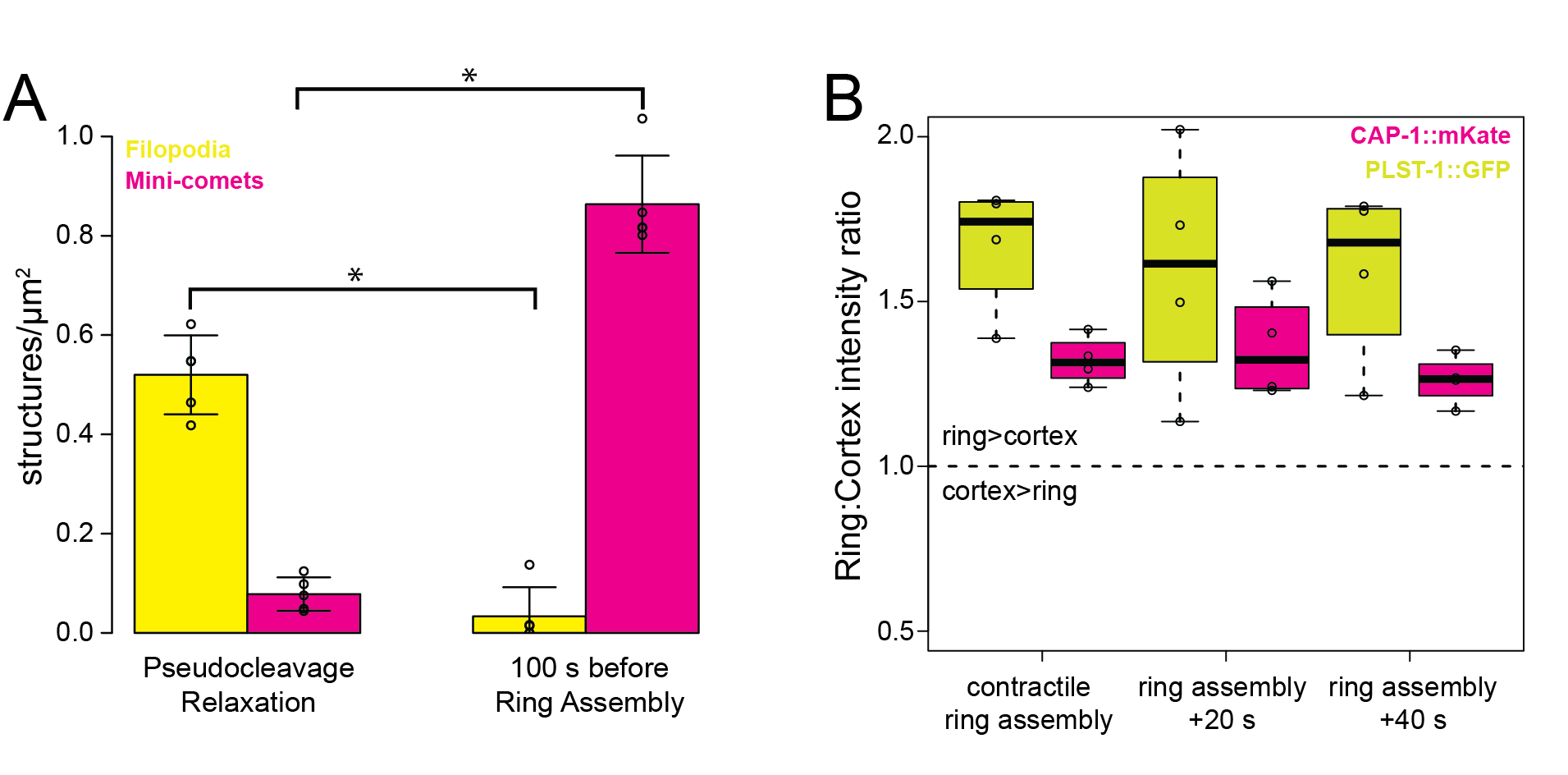

Figure S1. **Initial characterization of filopodia, mini-comets, and contractile ring, corresponding to Fig. 1. (A)** Number of filopodia and mini-comets at pseudocleavage relaxation and in late mitosis (100 s before cytokinesis onset) in embryos expressing LifeAct::mCherry. Filopodia assembly dominates during early mitosis, and mini-comet assembly dominates during late mitosis. For filopodia between the two time points, p=0.0002. For mini-comets, p=0.0001 (unpaired T-test). Error bars are standard deviation. **(B)** Enrichment of plastin PLST-1::GFP and CP CAP-1::mKate in the contractile ring during cytokinesis. Ratio of fluorescence intensities of a 35x100 pixel rectangular ROI spanning the contractile ring compared to an ROI of the same size on the posterior cortex of the embryo at three timepoints, from onset of contractile ring assembly to 40 s later. N=4 embryos.

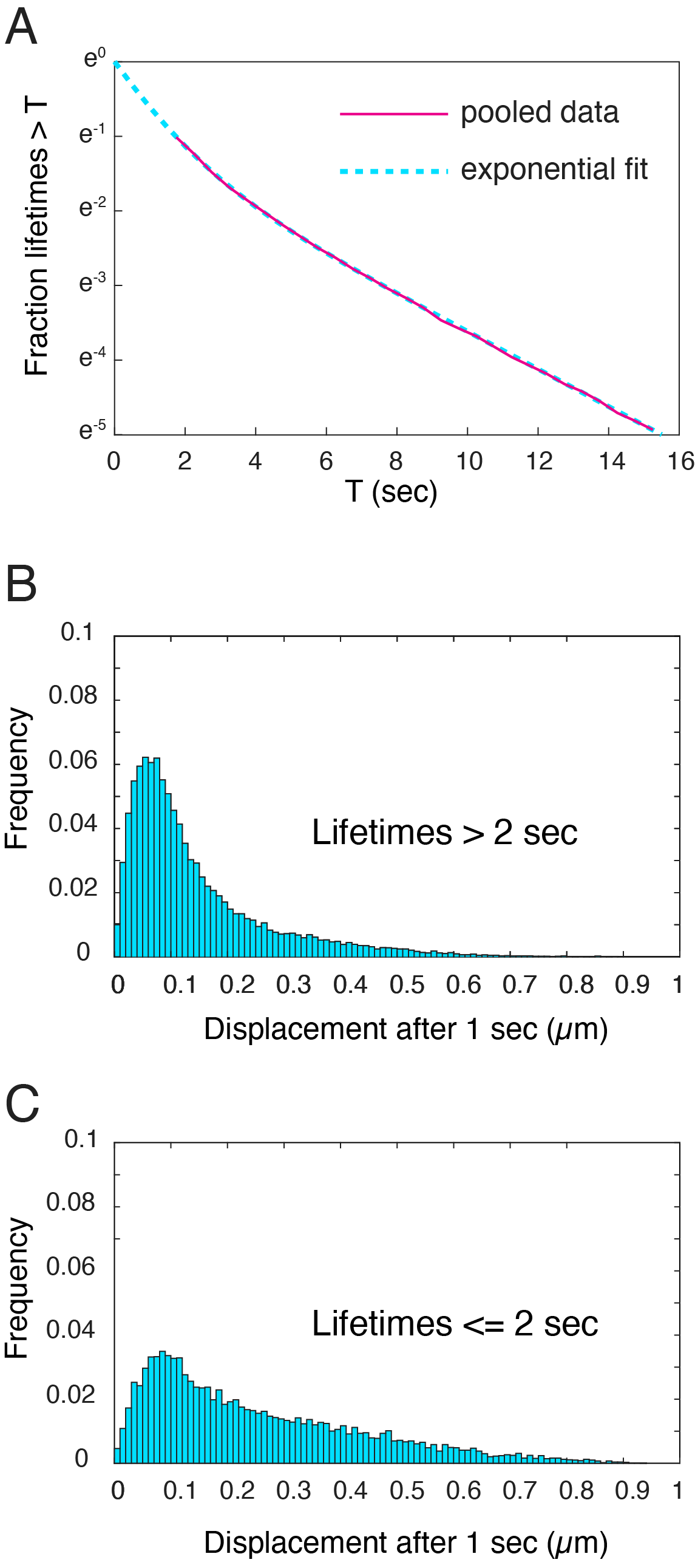

Figure S2. **Analysis of CAP-1::GFP cortex residence times *in vivo,* corresponding to Fig. 2 G-H. (A)** Distribution of CAP-1::GFP lifetimes at the cortex. Pink line: pooled average from N=9 embryos. Blue dotted line: non-linear least squares fit of the pooled data to a sum of two exponentials, consistent with a mixture and fast and slow dissociation kinetics. **(B-C)** Distribution of short-term (1 s) displacements of tracked CAP-1::GFP molecules with lifetimes **(B)** greater than 2 s, and **(C)** less than or equal to 2 s. Single molecules with lifetimes greater than 2 s have lower mobilities, suggesting that slow dissociation is associated with cortex binding. Thus, we used the slower of the two inferred dissociation rates as an estimate of the cortex dissociation rate.

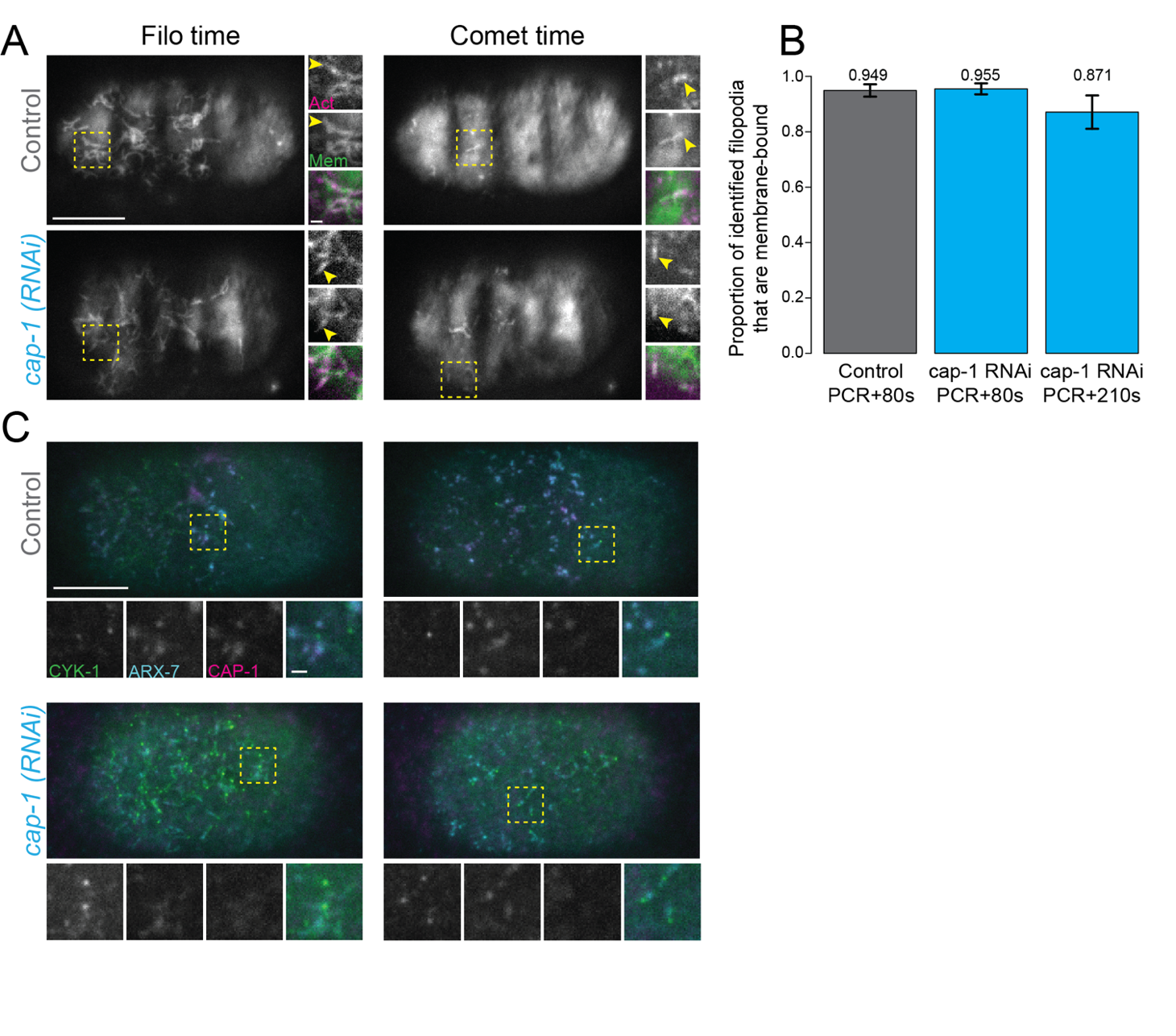
 Figure S3. **CeCP knockdown-induced ectopic filopodia contain the same components as control filopodia, corresponding to Fig. 3. (A)** Representative images of control (upper) and *cap-1(RNAi)* (lower) embryos during early (PCR+80 s, left) and late (PCR+210 s, right) mitosis expressing LifeAct::mCherry and GFP::membrane (PLC∂). A representative area, outlined in dashed yellow box, is shown in further detail to the right of each image. Yellow arrowhead points to a filopodium shaft. **(B)** Proportion of filopodia identified in LifeAct channel that colocalized with a membrane enrichment in control (gray) and *cap-1(RNAi)* (Blue) embryos. p>0.05 (n.s.), N=3 embryos for each condition. **(C)** Representative images of control (upper) and *cap-1(RNAi)* (lower) embryos during early (PCR+80 s, left) and late (PCR+210 s, right) expressing formin CYK-1::GFP, Arp2/3 complex component ARX-7::HALO and *Ce*CP CAP-1::mKate. Each yellow dashed box highlights a single filopodium which is enlarged and shown as separate channels, followed by merged image, below each image.

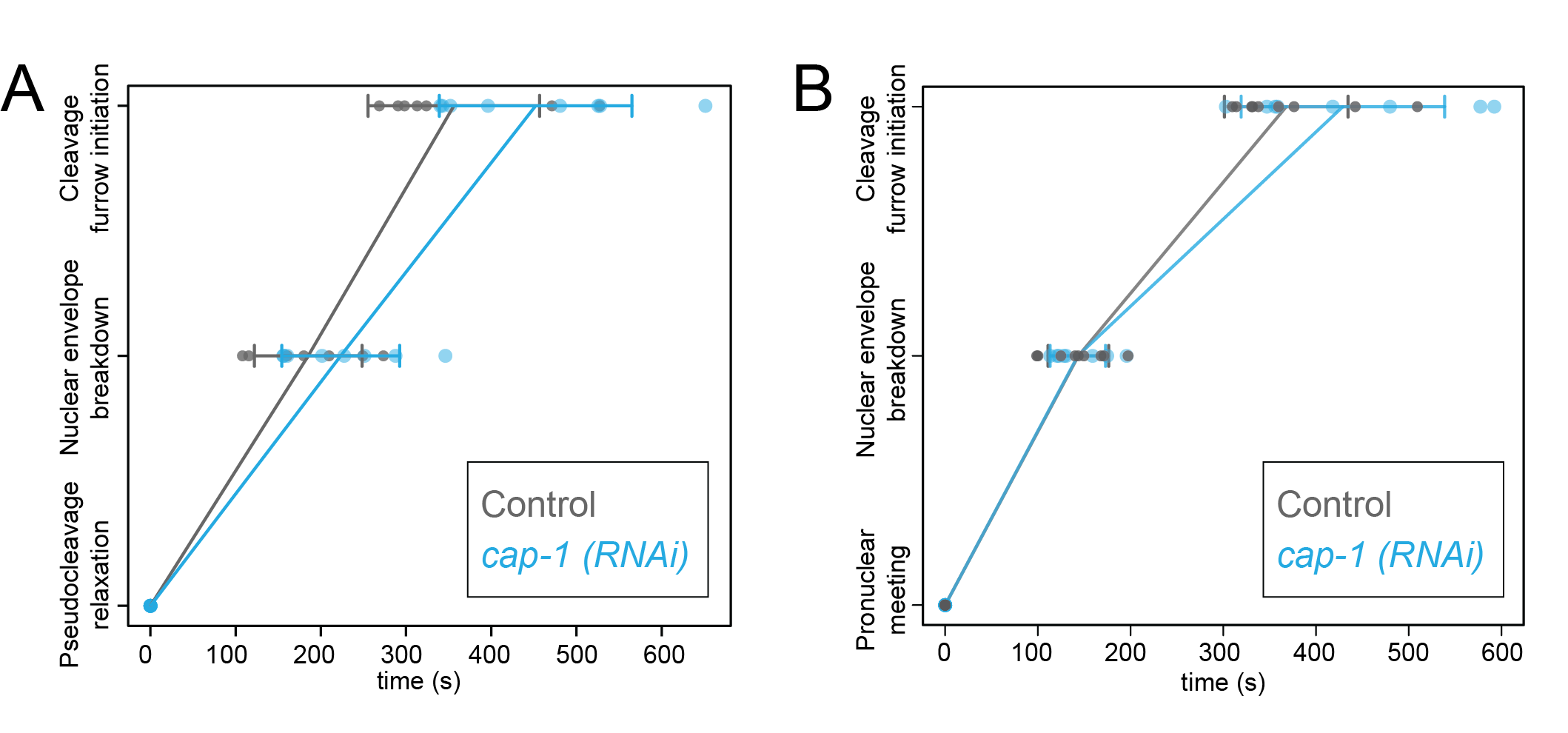

Figure S4**. CP KD does not significantly change progression of the cell cycle, corresponding to Fig. 3.** Quantification of cell cycle timing for control (gray) and cap-1(RNAi)-treated embryos (blue), using pronuclear position as determined by DIC microscopy. **(A)** Relative to pseudocleavage relaxation as t=0. **(B)** Relative to pronuclear meeting as t=0. Each dot represents a single embryo, N=8 embryos per condition. Error bars are standard deviation. P>0.05 (n.s.) for all timepoints measured, student’s T-test

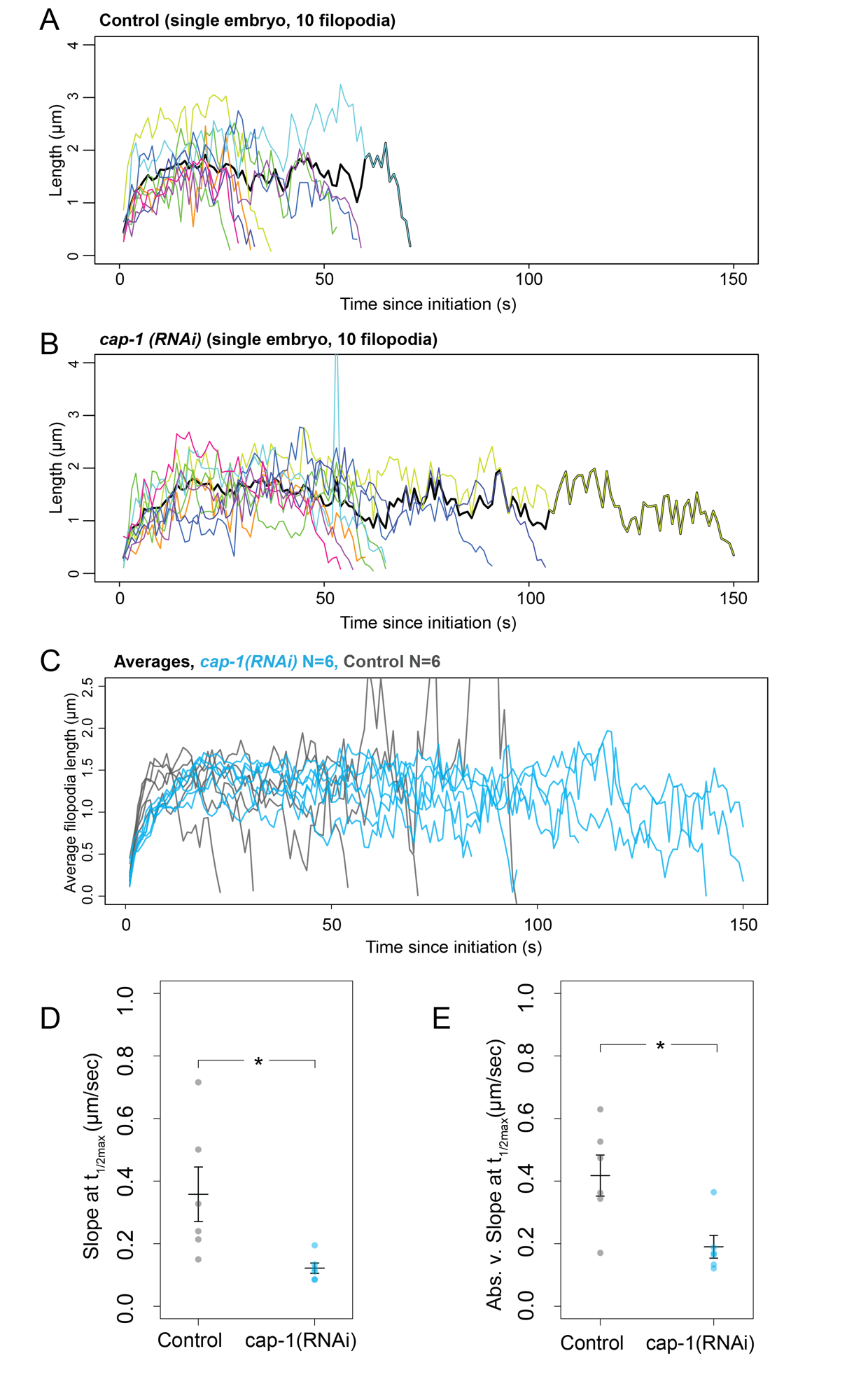

Figure S5. **Capping protein depletion changes filopodia assembly and disassembly dynamics, corresponding to Fig 5. (A-B)**. Length of 10 individual filopodia (thin colored lines) in a single representative embryo over the lifetime of each filopodium. Thick black trace represents the average filopodia length over time, as shown in C and Fig. 5 F and H. **(A)** Representative control embryo. **(B)** Representative cap-1(RNAi)-treated embryo. **(C)** Average filopodia lengths over time for control (gray) and *cap-1(RNAi)* (blue) embryos. Each line represents the average of 10 filopodia from a single embryo. N=6 for both conditions. **(D-E)** Slope at t_1/2max_ for filopodia assembly **(D)** and disassembly **(E).** Each dot represents the average of 10 filopodia in a single embryo. N=6 embryos for each condition. Mean of means +/- S.E.M., p=0.042 for assembly, p=0.017 for disassembly, unpaired t-test.

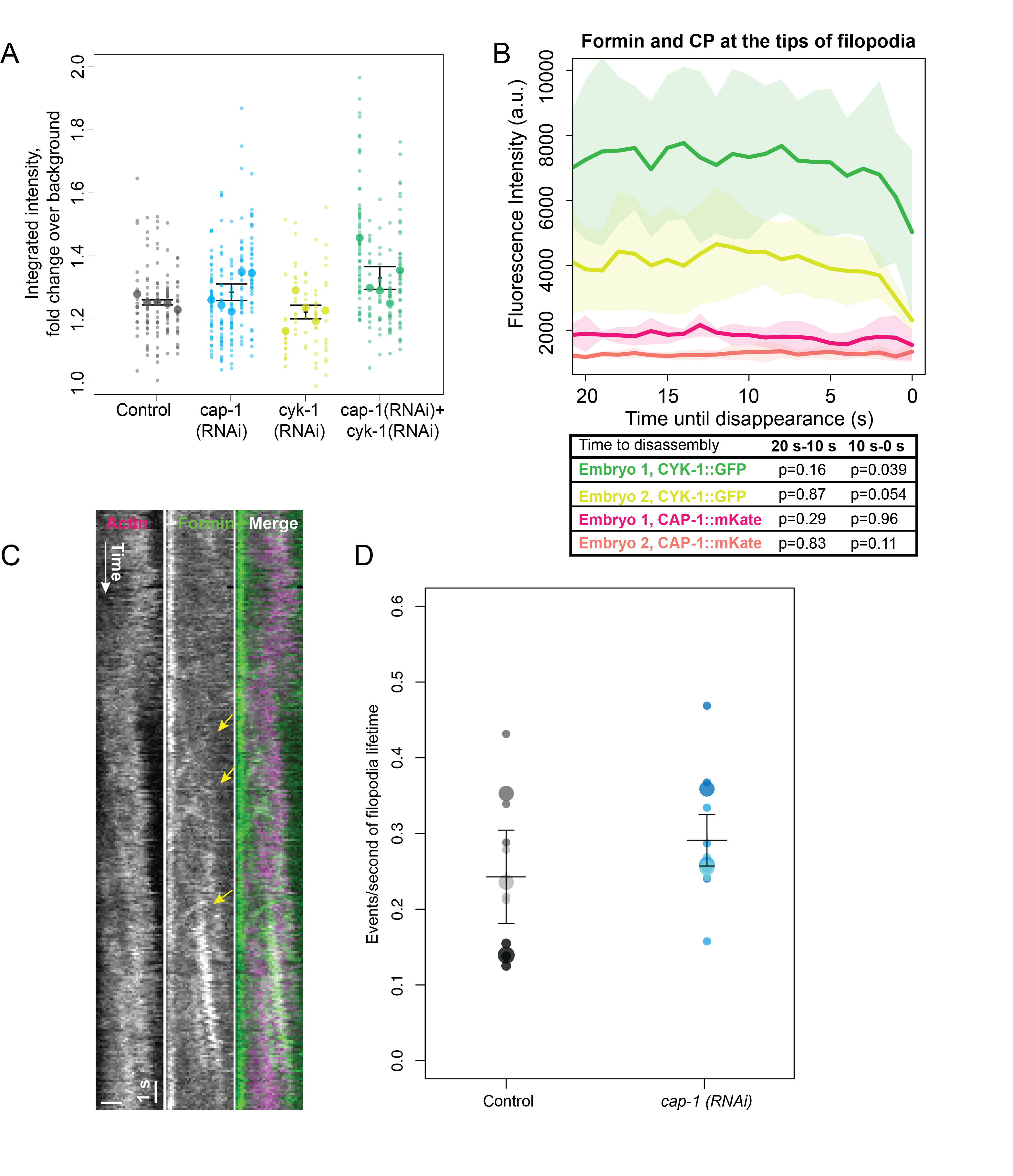

**Figure S6. Formin and capping protein compete in filopodia, corresponding to Fig. 6. (A)** Integrated intensity of filopodia cross sections, expressed as a fold increase over local background of each filopodium. Large dots represent the average of each embryo (N=5), small dots represent each individual filopodium in that embryo. Error bars: mean of means, +/- S.E.M. **(B)** Intensity of CYK-1::GFP and CAP-1::mKate at the tips of filopodia in the last 20 seconds of filopodia lifetime. Each line is an average of 5 filopodia, shaded areas are standard deviation. N=2 embryos. (lower) Table comparing change over time in values from above graph. Formin intensity decreases in the last 10 seconds, CAP-1 intensity does not change. **(C)** Kymograph of a single representative filopodium in a control-treated embryo expressing LifeAct::mCherry and formin CYK-1::GFP. Filopodium is aligned with the tip to the left, base to the right. Yellow arrowheads point to events counted as fast anterograde formin movements. **(D)** Frequency of fast anterograde formin events in control (gray) and *cap-1(RNAi)*-treated (blue) embryos. Small dots: n=3 filopodia per embryo, large dots: N=3 embryos per condition.

**Video legends**

Video 1. **Whole-cell imaging of plastin and capping protein, corresponding to Fig. 1 B.** Near-TIRFM imaging of a *C. elegans* zygote expressing endogenously tagged plastin PLST-1::GFP (yellow) and capping protein subunit CAP-1::mKate (magenta) undergoing the first cell cycle of development. Circles labeled ‘C’ and ‘D’ correspond to filopodia and mini-comets from Fig. 1 C and D respectively. Time 0 is onset of pseudocleavage relaxation. Scalebar is 10 µm. Frames collected at 1 fps, displayed at 50 fps.

Video 2. **Insets of filopodia and mini-comets, corresponding to Fig. 1 C-D.** Near-TIRFM imaging of a *C. elegans* zygote expressing endogenously tagged plastin PLST-1::GFP (yellow) and capping protein subunit CAP-1::mKate (magenta), insets from Video 1. Upper row depicts a representative filopodium during early mitosis. Filled arrowhead points to CAP-1::mKate enrichment at the base of the filopodium. Lower row depicts representative mini-comets during late mitosis. Unfilled arrowhead points to CAP-1::mKate enrichment in mini-comets. Frames collected at 1 fps, displayed at 10 fps.

Video 3. **Whole-cell imaging of formin, Arp2/3 complex, and capping protein, corresponding to Fig. 1 E.** Near-TIRFM imaging of a *C. elegans* zygote expressing endogenously tagged plastin ARX-1::HALO (cyan), formin CYK-1::GFP (green), and capping protein subunit CAP-1::mKate (magenta) undergoing the first cell division of development. Circles labeled ‘E ii’ and ‘E iv’ correspond to filopodium and mini-comet from Fig. 1 E ii and iv respectively. Time 0 is onset of pseudocleavage relaxation. Scalebar is 10 µm. Frames collected at 1 fps, displayed at 40 fps.

Video 4. **Insets of filopodium and mini-comets, corresponding to Fig. 1 E.** Near-TIRFM imaging of a *C. elegans* zygote expressing endogenously tagged plastin ARX-1::HALO (cyan), formin CYK-1::GFP (green), and capping protein subunit CAP-1::mKate (magenta), insets from Video 3. (Upper) A representative filopodium during early mitosis. Unfilled arrowhead indicates CYK-1::GFP-rich filopodia tip. Filled arrowhead indicates CAP-1::mKate enrichment in ARX-7::HALO-rich base. (Lower) Representative mini-comet during late mitosis. Unfilled arrowhead indicates CAP-1::mKate enrichment in ARX-7::HALO-rich mini-comet. Frames collected at 1 fps, displayed at 20 fps.

Video 5. **Single filament imaging of F-actin and capping protein *in vitro,* corresponding to Fig. 2 E.** Two-color TIRFM visualization of the assembly of 1.5 µM Mg-ATP *C. elegans* actin (10% alexa 488-labeled) (yellow) with 647-labeled CP (magenta). Representative image of *Ce*CP-647 (magenta arrowhead) at the barbed end of an actin filament. Magenta arrowhead indicates CP binding event. Collected at 0.2 fps, displayed at 10 fps. Scalebar is 5 µm.

Video 6. **Single molecule imaging of CAP-1::GFP, corresponding to Fig. 2 G.** Near-TIRFM imaging of representative zygote expressing CAP-1::GFP as a transgene. Imaged in late mitosis, after 24 h of *GFP(RNAi).* Yellow circle: a single representative CP-cortex binding event. Imaged at 2 fps, displayed at 10 fps.

Video 7. **Whole-cell imaging of F-actin during mitosis upon control and *cap-1(RNAi)* treatment, corresponding to Fig. 3 A*.*** Near-TIRFM imaging of representative L4440 control-treated (left) and 46 hours of L4440 *cap-1(RNAi)-treated* (right) zygotes expressing LifeAct::mCherry. Time 0 is onset of pseudocleavage relaxation. Collected at 1 fps, displayed at 40 fps.

Video 8. **Insets of representative filopodium and mini-comets from Video 7 (control), corresponding to Fig. 3A.** Near-TIRFM images of control zygote expressing LifeAct::mCherry. (Left) Representative filopodium, where unfilled arrowhead indicates the tip, filled arrowhead indicates the base. (Right) representative mini-comet, indicated by arrowhead. Scalebar is 1 µm. Collected at 2 fps, displayed at 20 fps.

Video 9. **Whole-cell imaging of F-actin and membrane, corresponding to Fig. S3 A.** Representative near-TIRFM images of control (upper) and 24 hours of T444T *cap-1(RNAi*)-treated (lower) zygotes expressing LifeAct::mCherry (magenta); GFP::membrane (PLC∂) (green). Collected at 0.5 fps, displayed at 25 fps.

Video 10. **Whole cell imaging of F-actin upon various *RNAi*-treatments, corresponding to Fig. 6 A.** Each image is a representative near-TIRFM image of a zygote of the indicated treatments. Time 0 is onset of pseudocleavage relaxation. Scalebar is 10 µm. Collected at 1 fps, displayed at 50 fps.

Video 11. **Whole-cell fast imaging of F-actin and formin, corresponding to Fig. S5 C.** Representative near-TIRFM image of control-treated zygote expressing LifeAct::mCherry (magenta) and endogenously tagged formin CYK-1::GFP (green). Collected at 10 fps, displayed at 100 fps.

Video 12. **Inset of representative filopodium from Video 11, corresponding to Fig. S5 C.** Representative near-TIRFM fast imaging of a filopodium from a zygote expressing LifeAct::mCherry (magenta) and endogenously tagged formin CYK-1::GFP (green). Collected at 10 fps, displayed at 100 fps.

**Table 1. Worm and RNAi strains used**

| **Reagent** | **Source** | **Info** |
| --- | --- | --- |
| **Tagged genes** |  |  |
| PLST-1::GFP | Ronen Zaidel-Bar (Ding et al <https://doi.org/10.1083/jcb.201603070>) | plst-1(msn190[plst-1::gfp]) IV |
| CAP-1::mKate | Ronen Zaidel-Bar (Ray et al <https://doi.org/10.1242/dev.201099>) | cap-1 (knu650 [pNU1550 - N-terminal degron mKate2 unc119(+)] |
| CYK-1::GFP | Ronen Zaidel-Bar (Padmanabhan et al <https://doi.org/10.1016/j.cub.2016.10.032>) | cyk-1 (knu83 C-terminal GFP, unc-119 (+)) |
| LifeAct::mCherry^1^ | Chris Pohl, Zhirong Bao  ^1.BV70 annotated on CGC as LifeAct::RFP^ | zbIs2[pie-1::Lifeact::mCherry, unc-119(+)] |
| ARX-7::HALO | Munro/Kovar Lab (Rachel Kadzik, SunyBiotech, this study) | Arx-7::HALO I |
| Membrane::GFP | Goldstein Lab (Heppert et al [10.1091/mbc.E16-01-0063](https://doi.org/10.1091/mbc.E16-01-0063)) | cpIs53 [mex-5p::GFP-C1::PLC(delta)-PH::tbb-2 3'UTR + unc-119 (+)] II. |
| CAP-1::GFP | Ronen Zaidel-Bar (Nemametrics, COP2261/COP2262) | knuSi859 [pNU2577(RZAI65 sip-1p::cap-1::GFP::tbb-2 3'UTR insertion in ttTi5605, unc-119(+))] II; unc-119(ed3) III |
| **Crossed worm strains** |  |  |
| PLST-1::GFP; CAP-1::mKate | Ronen Zaidel-Bar |  |
| LifeAct::mCherry; CYK-1::GFP | Munro Lab |  |
| LifeAct::mCherry; Membrane::GFP | Rachel Kadzik |  |
| CYK-1::GFP; ARX-7::HALO; CAP-1::mKate | Sarah Yde |  |
| **RNAi** |  |  |
| L4440 control | Fire lab, Addgene plasmid # 1654 ; http://n2t.net/addgene:1654 ; RRID:Addgene_1654 | Empty vector control |
| CAP-1 L4440 | Ahringer feeding library (Kamath et al https://doi.org/10.1038/nature01278) | D2024.6 in L4440 vector |
| CYK-1 L4440 | Ahringer feeding library (Kamath et al https://doi.org/10.1038/nature01278) | F11H8.4 in L4440 vector |
| T444T control | Vellai Lab, Addgene plasmid # 113081 ; http://n2t.net/addgene:113081 ; RRID:Addgene_113081 | Empty vector control |
| CAP-1 T444T | Zaidel-Bar lab | D2024.6 (NT 208-611) in T444T vector |
| GFP | Jeremy Nance, generated by the Fire lab and available at <http://www.addgene.org/1649/> | L4417 vector |
